# A systematic characterization of intrinsically formed microglia-like cells during retinal organoid differentiation

**DOI:** 10.1101/2021.12.30.474511

**Authors:** Katarina Bartalska, Verena Hübschmann, Medina Korkut-Demirbaş, Ryan John A. Cubero, Alessandro Venturino, Karl Rössler, Thomas Czech, Sandra Siegert

**Author notes:** Contributed equally (Katarina Bartalska, Verena Hübschmann, Medina Korkut-Demirbaş).

## Abstract

Brain organoids differentiated from human induced pluripotent stem cells provide a unique opportunity to investigate the development, organization and connectivity of neurons in a complex cellular environment. However, organoids usually lack microglia, brain-resident immune cells which are both present in the early human embryonic brain and participate in neuronal circuit development.

Here, we find that microglia innately develop in unguided retinal organoid differentiation between week 3 and 4 in 2.5D culture and appear later in floating, non-pigmented, 3D-cystic compartments. We enriched for cystic structures using a low-dosed BMP4 application and performed mass spectrometry, thus defining the protein composition of microglia-containing compartments. We found that cystic compartments expressed both mesenchymal and epithelial markers with microglia enriched in the mesenchymal region. Interestingly, microglia-like cells started to express the border-associated macrophage marker CD163. The preferential localization of human microglia to a mesenchymal compartment provides insight into the behavior and migration of microglia. The model will ultimately allow detailed study of these enigmatic cells and how they enter and distribute within the human brain.

## Introduction

The human brain consists of about 80 billion neurons, 60 billion glial cells and 20 billion endothelial cells that self-organize during development into cellular networks that perform distinct functions (Barresi, 2020). Our current knowledge about how these cells form and connect in the human brain is limited to observations from post-mortem fetal brain studies or reports from model organisms including mice. Microglia, the brain parenchymal immune cells, fine-tune neuronal circuits at the cellular- and synaptic level (Cunningham et al., 2013; Guizzetti et al., 2014; Paolicelli and Gross, 2011; Schafer et al., 2012; Squarzoni et al., 2014). They derive from a primitive macrophage population, which develops within the yolk sac in both mouse (Ginhoux et al., 2010) and humans (Bian et al., 2020) and therefore represent a distinct macrophage population as they occur prior to the onset of hepatic and bone marrow hematopoiesis (Juul and Christensen, 2018; Menassa and Gomez-Nicola, 2018).

Immunostaining of human embryonic brain tissue indicate that microglia enter the cerebral wall from the ventricular lumen and the leptomeninges at 4.5 gestational weeks and gradually colonize the cortex (Monier et al., 2007; Rezaie et al., 2005). The critical role of microglia in early human brain development has been further supported by hereditary mutations in macrophage-selective genes that cause numerous structural brain malformations in pediatric leukoencephalopathy (Oosterhof et al., 2019). A current bottleneck is the lack of accessible models that accurately recapitulate human microglia development, distribution and action during circuit formation.

Human induced pluripotent stem cells (hiPSC) have revolutionized the field of tissue engineering and allowed exploration of aspects of embryonic brain development (Bagley et al., 2017; Camp et al., 2015; Cowan et al., 2020; Lancaster et al., 2017). However, mesoderm-derived microglia are commonly lacking within brain organoids (Collin et al., 2019; Cowan et al., 2020; Ginhoux et al., 2010; Kim et al., 2019; Lancaster and Knoblich, 2014). One likely explanation for this is that differentiation protocols often use supplements to direct hiPSC-formed embryoid bodies (EB) toward one of the three germ layers, e.g., brain organoids toward the neuroectodermal lineage (Chambers et al., 2009; Lancaster et al., 2017; Tanaka and Park, 2021). In attempts to obtain human microglia, guided protocols were established to enrich for them (Abud et al., 2017; Guttikonda et al., 2021; Haenseler et al., 2017; McQuade et al., 2018; Muffat et al., 2016; Pandya et al., 2017; Takata et al., 2017; 2011). To analyze microglia function and interaction with neurons, several groups have combined these microglia-like cells with separately-derived cerebral organoids (Abud et al., 2017; Song et al., 2019; Xu et al., 2021). A major drawback is that this assembly strategy does not capture the natural progression of microglial appearance and distribution within brain tissue.

Unguided brain organoid differentiation provides an alternative methodology in which EBs are cultured with minimal external interference based on intrinsic self-organization principles leading to a variety of cell lineage identities from fore-, mid- and hindbrain (Qian et al., 2019). One of the first brain region-specific protocols focuses on retinal organoids (Eiraku et al., 2011; Nakano et al., 2012). The method reliably recapitulates the typical eye cup structure, expresses markers of well-defined cell types, and shows a light-sensitive response (Cowan et al., 2020; Zhong et al., 2014). The data regarding intrinsic microglia occurrence are controversial. Whereas single-cell RNA-sequencing of brain organoids with bilateral optic vesicles identified a glial cluster expressing microglia-specific markers (Gabriel et al., 2021), other studies do not report, or the provided data do not support, their presence in retinal organoids (Collin et al., 2019; Cowan et al., 2020; Kim et al., 2019). A microglial transcriptional signature can be detected in human embryonic retinal tissue by gestational week 5 (Hu et al., 2019; Mellough et al., 2019) suggesting that microglia appear early in development. Microglia are localized within the human retinal layers by gestational week 10 and frequently show a phagocytic phenotype (Diaz-Araya et al., 1995). To clarify whether microglia develop in unguided retinal organoid-focused differentiation and are present during retinal development, we implemented the protocol from (Zhong et al., 2014). We stained 2.5D- and 3D cultures with the pan-macrophage marker IBA1/AIF1 (ionized calcium binding adaptor molecule 1/ allograft inflammatory factor 1), which identifies brain parenchymal- (microglia), blood-derived- (MΦ), and border-associated- (perivascular pvMΦ, leptomeningeal mMΦ, choroid plexus cpMΦ) macrophages (Imai et al., 1996; Ito et al., 2001; Kierdorf et al., 2019; Prinz and Priller, 2014).

Here, we consistently find that IBA1^+^-cells can be detected in 2.5D culture by differentiation week 3 to 4 in parallel to developing retinal organoids. We confirmed that these represent microglia-like cells using specific marker expression. Microglia preferentially occupied non-pigmented, cystic compartments in 2.5D and 3D cultures, structures that are commonly overlooked in organoid-focused studies. To further analyze these cystic structures, we enriched for them by applying BMP4 at a low-dose. The cystic compartments showed high expression of mesenchymal and epithelial markers Vimentin and E-Cadherin, respectively. We confirmed a similar expression pattern in the cystic compartments in retinal organoids grown without BMP4. Since a mesenchymal/epithelial environment is typical of the developing lymphatic and circulatory system (MacCord, 2012; Pill et al., 2015; Wimmer et al., 2019), we stained the IBA1^+^-cells for the border-associated marker CD163. Indeed, we found strong overlap between IBA1 and CD163 expression, the latter of which is turned on between week 5 and 6 in 2.5D culture.

In summary, our results confirm that IBA1^+^-cells intrinsically develop in retinal organoid cultures and map their presence to cystic mesenchymal-like regions which co-develop next to 3D-retinal organoids during differentiation. This work offers a model for exploring microglia integration during early development, and provides a foundation for future studies to dissect the molecular signaling mechanisms that attract microglia and foster their incorporation into brain organoids.

## Results

### IBA1^+^-microglia-like cells appear during retinal organoid differentiation

To identify whether 3D-retinal organoids contain intrinsic microglia-like cells, we applied an established retinal organoid-differentiation protocol (Zhong et al., 2014) to two hiPSC lines of different origins (**Fig. 1a, Supplementary Fig. 1a-b**). One hiPSC line was originally derived from a 60^+^-old skin fibroblast donor (SC102A) and the other from fetal umbilical cord blood cells (CR05, **Supplementary Table 1**). Both hiPSC lines behaved similarly and formed typical optic-cup structures within four weeks in 2.5D culture. They developed further into anatomically comparable 3D-retinal cups (**Supplementary Fig. 1c**) expressing cell type-specific markers for photoreceptor-, bipolar-, amacrine-, ganglion and Müller glial cells by week 18 (**Supplementary Fig. 1d**) (Hoshino et al., 2017; Luo et al., 2019; Zhang et al., 2019). We used IBA1 as a marker for microglia and confirmed the antibody functionality in human temporal lobe brain tissue, where IBA1 labeled parenchymal microglia and pvMΦ (**Supplementary Fig. 2**). When we immunostained our 3D-retinal organoids, we commonly observed no cell-defined IBA1 staining (**Fig. 1b**). Occasionally, we found a few IBA1^+^-cells close to the retinal-cup (**Supplementary Fig. 3a**) but the cells were not numerous or as deeply integrated as described for the human embryo retina at similar age (Diaz-Araya et al., 1995).

**Figure 1.**
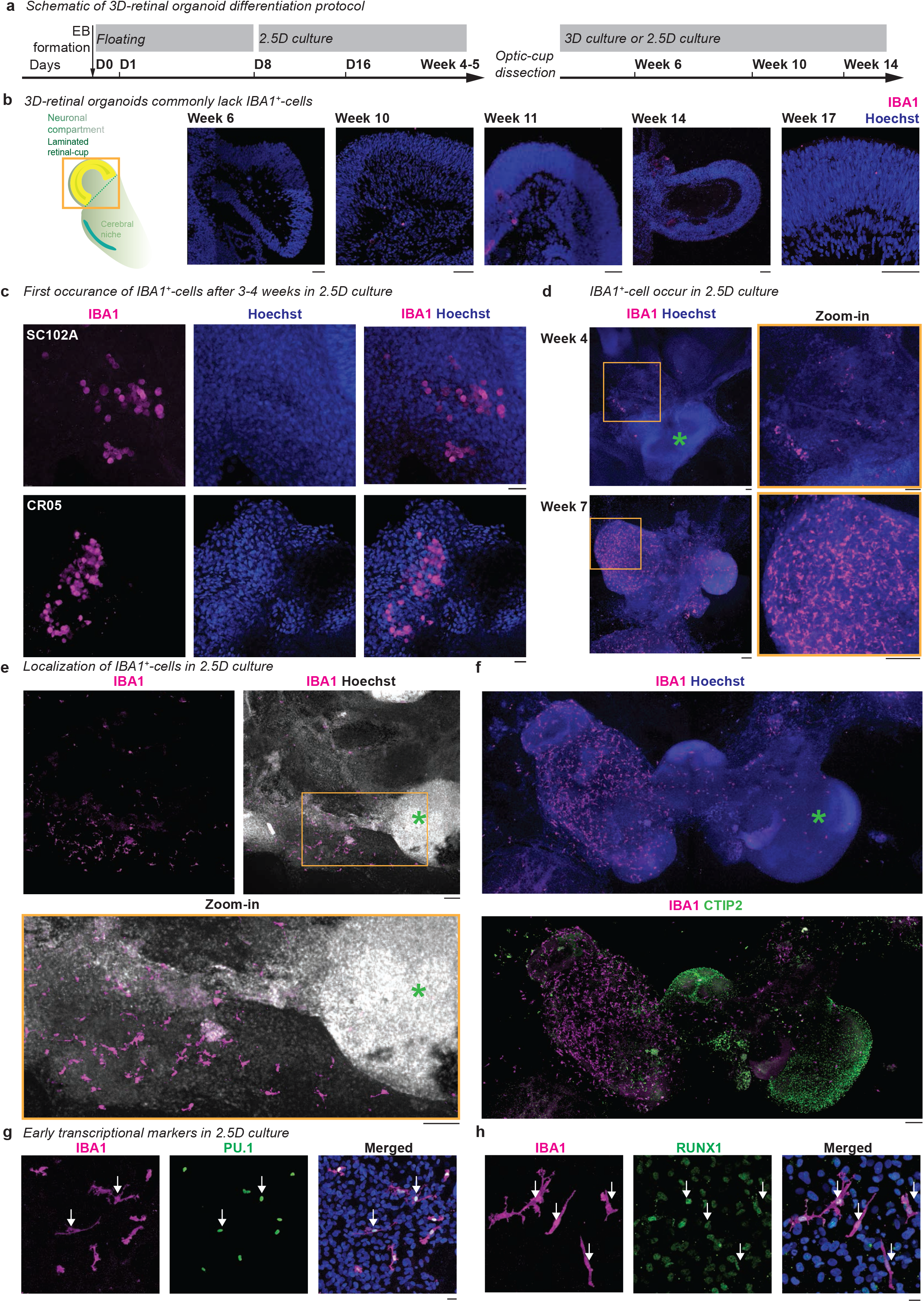
IBA1^+^-microglia-like cells occur during retinal organoid differentiation. **a**, Schematic of retinal organoid differentiation protocol (see also **Supplementary Fig. 1a** for detailed information about media composition). D, days after induced differentiation. EB, embryoid bodies. **b**-**h**, Immunostaining for IBA1 (ionized calcium-binding adapter molecule 1, magenta) with Hoechst to highlight nuclei (blue, except white in **e**). **b**, Cryostat section of 3D-retinal organoids with focus on retinal-cup at week 6, 10, 11, 14, and 17 for SC102A. Note: IBA1 staining occasionally occurred as a layered or dotted structure, which did not resolve in distinct cell morphologies. We excluded such staining patterns from further interpretations. Scale bar: 50 μm. **c**-**h**, Staining in 2.5D culture. **c**, First occurrence of IBA1^+^-cells in SC102A (top) and CR05 (bottom) between week 3 and 4. Scale bar: 20 μm. **d**, SC102A at week 4 (top) and 7 (bottom). Scale bar: 100 μm. **e**, SC102A at week 9. Scale bar: 100 μm. **f**, SC102A at week 7. Green *, cell-dense area. Immunostaining for CTIP2 (COUP-TF-Interacting-Protein 2, green). Scale bar: 150 μm. **g**-**h**, SC102A at week 5, immunostained in green: **g**, PU.1 (hematopoietic transcription factor PU.1). **h**, RUNX1 (runt-related transcription factor 1). White arrow, overlap. Scale bar: 20 μm.

Based on this rare presence of microglia, we hypothesized that IBA1^+^-cells might be enriched in a compartment other than the retinal cup. Thus, we revisited the 2.5D culture, prior to dissection of optic-cups at week 4 (**Fig. 1a**). Between week 3 and 4, we found clusters of IBA1^+^-cells (**Fig. 1c**), which start to spread within the culture by week 4 and occupy distinct compartments by week 7 (**Fig. 1d**). These compartments were commonly less nuclei-dense (**Fig. 1e**). To investigate whether these compartments contained cortical cell types, we stained the 2.5D culture with CTIP2/BCL11b (BAF chromatin remodeling complex subunit), a marker expressed in the neocortex from early embryonic stages (Qian et al., 2016). Remarkably, the majority of IBA1^+^-cells were distinct from the CTIP2^+^-region (**Fig. 1f**), and if they were present, they mostly localized to the surface of these structures. To confirm that IBA1^+^-cells were microglia-like, we immunostained the 2.5D cultures between week 4 and 5 for the hematopoietic lineage-specific markers RUNX1 (Ginhoux et al., 2010), PU.1 (Kierdorf et al., 2013), and MYB (Schulz et al., 2012). As expected, all IBA1^+^-cells were positive for RUNX1 and PU.1 and negative for MYB (**Fig. 1g-h, Supplementary Fig. 3c-d**). The IBA1^+^-cells also expressed the mononucleate hematopoietic cell marker CD45 (Monier et al., 2007) (**Supplementary Fig. 3e**), the fractalkine receptor CX3CR1 (Hulshof et al., 2003) (**Supplementary Fig. 3f**), and the purinergic receptor P2Y12 (Mildner et al., 2017) (**Supplementary Fig. 3g**). Importantly, all markers were cross-validated for their specificity in human brain tissue (**Supplementary Fig. 2**). The IBA1^+^-cells were morphologically branched and frequently presented phagocytic cups (**Supplementary Fig. 3b**). Half of the IBA1^+^-cells co-expressed KI-67 indicating that they are in a proliferative state (Gerdes et al., 1984) (**Supplementary Fig. 3h**). The cells also expressed the mitotic marker phosphorylated histone H3 (PHH3) (Hirata et al., 2004) (**Supplementary Fig. 3i**). This characterization suggests that IBA1^+^-cells represent microglia-like cells that emerge within the retinal organoid differentiation protocol by week 4 and, notably, do not extensively populate retinal or cortical regions.

### IBA1^+^-cells enrich in defined cystic compartments

The presence of IBA1^+^-cells in less nuclear-dense structures inspired us to revisit our 3D-floating retinal “aggregates” (Cowan et al., 2020; Zhong et al., 2014) (**Fig. 2a**). We identified different structures, which we classified into three categories: first, 3D-retinal organoids that usually contained retinal-cup and cerebral regions (**Fig. 2a**). These regions can be characterized with OTX2 (Orthodenticle Homeobox 2) (Hoshino et al., 2017) and CTIP2, respectively, and were clearly distinct at week 9 (**Fig. 2b**). Second, we described 3D-cysts, which were semi-transparent, contained various-sized lumens and occasionally developed pigmentation or a cuboidal-shaped epithelial surface (**Fig. 2a**). Third, we identified 3D-retinal organoids in which a cystic compartment intrinsically co-developed (3D-retinal organoid-attached cyst, **Fig. 2a**). Interestingly, we found that IBA1^+^-cells were enriched in the cystic compartment of both 3D-cysts (**Fig. 2c**) and 3D-retinal organoids-attached cysts (**Fig. 2d**). Microglia-like cells distribute within the cystic compartment and are rarely associated with the surface cell layer. Cystic compartment that was pigmented tended to lack IBA1^+^-cells (**Fig. 2e**). This suggests that intrinsically-generated IBA1^+^-cells during retinal organoid differentiation preferentially occupy non-pigmented cystic compartments.

**Figure 2.**
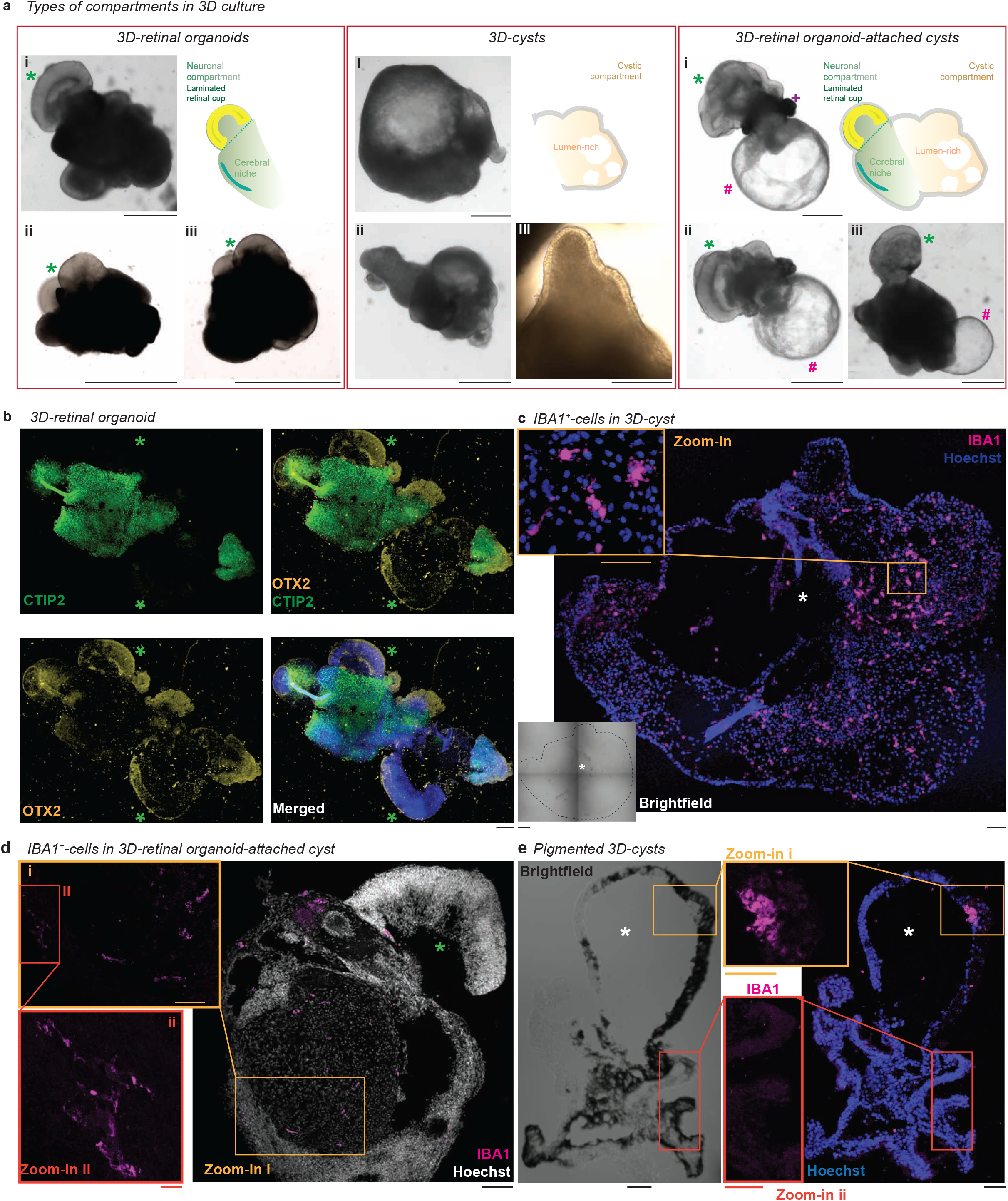
IBA1^+^-cells occupy 3D-cystic compartments. **a**, Brightfield images of different structures occurring within 3D-retinal organoid differentiation at week 5 for SC102A. Three representative images (i – iii) for each structure are depicted. Green *, retinal-cup. Magenta #, cystic compartment. Purple +, pigmented region in 3D-retinal organoid-attached cyst (i). Scale bar: 1000 μm, except 3D-cysts (iii) focusing on an epithelial-like structure: 200 μm. **b**-**e**, Immunostaining of cryostat sections for IBA1 (ionized calcium-binding adapter molecule 1, magenta), nuclei stained with Hoechst (blue, except **d** in white). **b**, 3D-retinal organoid (week 8-9) from SC102A stained for OTX2 (orthodenticle homeobox 2, orange) and CTIP2 (COUP-TF-Interacting-Protein 2, green). Green *, retinal cup. Scale bar: 150 μm. **c**, 3D-cyst from SC102A (week 8-9) with bright field image. Dashed-line, cyst surrounding. White *, cystic lumen. Scale bar: 100 μm. Zoom-in to IBA1^+^-cells. Scale bar: 50 μm. **d**, 3D-retinal organoid-attached cyst for SC102A (9-10 weeks). Scale bar: 100 μm. Zoom-in: Scale bar: 50 μm (i), 10 μm (ii). **e**, Pigmented 3D-cyst (week 8-9) from SC102A with bright field image. White *, cystic lumen. Scale bar: 100 μm, zoom-in: 50 μm.

### IBA1^+^-cells populate cystic compartments generated with BMP4-guided differentiation

Next, we sought insight into the tissue identity of IBA1^+^-cell enriched cystic compartment. However, we were confronted with the heterogeneity of our cultures: first, we saw variable numbers of generated cystic compartments upon differentiation and second, these structures were occupied with different numbers of IBA1^+^-cells. Thus, we looked for an alternative, more consistent strategy to generate 3D-cysts similar to our retinal organoid differentiation protocol. Recent hiPSC-derived microglia-like protocols have reported that bone morphogenetic protein 4 (BMP4) promotes microglia generation *in vitro* (Abud et al., 2017; Guttikonda et al., 2021; Haenseler et al., 2017; McQuade et al., 2018; Muffat et al., 2016; Pandya et al., 2017; Takata et al., 2017; 2011) and mentioned the development of cystic structures without providing additional details (Muffat et al., 2016; Vaughan-Jackson et al., 2021). In order to enrich for microglia in our cultures, we applied low-dosed BMP4 one day after EB formation to otherwise unchanged retinal organoid differentiation protocol (**Fig. 3a**). Optic-cup structures have no longer been formed in both hiPSC lines, instead, EBs formed an irregular shape, and after seeding, developed non-pigmented 3D-cysts that started to float by week 3 (**Fig. 3b**). These 3D-cysts were the only developing 3D-structure formed by BMP4-guided differentiation in both hiPSC lines (**Fig. 3b, Supplementary Fig. 4a**). From week 4 onwards, small branched cells started to float in the supernatant (**Fig. 3c**), which have previously been described as microglia-like cells (Haenseler et al., 2017). To verify this cell identity, we either collected and seeded these cells for immunostaining or directly labeled the 2.5D culture. The cells were IBA1^+^ and expressed RUNX1, PU.1, CD45, and P2Y12 (**Supplementary Fig. 4b-g**) suggesting that they are microglia-like cells similar to the non-BMP4-treated population (**Fig. 1g-h, Supplementary Fig. 3c-g**). To identify whether IBA1^+^-cells also occupied 3D-cysts, we collected cysts at several time points after differentiation and performed wholemount immunostaining. Starting at week 5.5, IBA1^+^-cells populated the 3D-cysts (**Fig. 4a**) and increased in number over time (**Fig. 4b**). Due to the appearance of IBA1^+^-cells in the supernatant and their occurrence in the 3D-cyst, we hypothesized that the 3D-cysts might be the source of IBA1^+^-cells. To test this, we collected 3D-cysts from a 2.5D culture plate at week 2.5, 3, 4, and 5 and cultured these separately, in parallel with the left-over 3D-cysts. These cultures were developed until week 7.5 and then immunostained for IBA1 (**Fig. 4c**). The 3D-cysts isolated at week 2.5, 3, and 4 unexpectedly contained only a few IBA1^+^-cells. In contrast, 3D-cysts isolated at week 5 had a similarly high number of IBA1^+^-cells to the 3D-cysts cultured on the original plate (**Fig. 4d**) excluding 3D-cysts as the source of IBA1^+^-cells. Therefore, we revisited the 2.5D culture and found that IBA1^+^-cells appeared on the plate between week 2 and 3 (**Fig. 4e**) suggesting that 3D-cysts are occupied by IBA1^+^-cells similar to those observed in the non-BMP4 treated retinal organoid differentiation in 2.5D culture (**Fig. 1c-f**).

**Figure 3.**
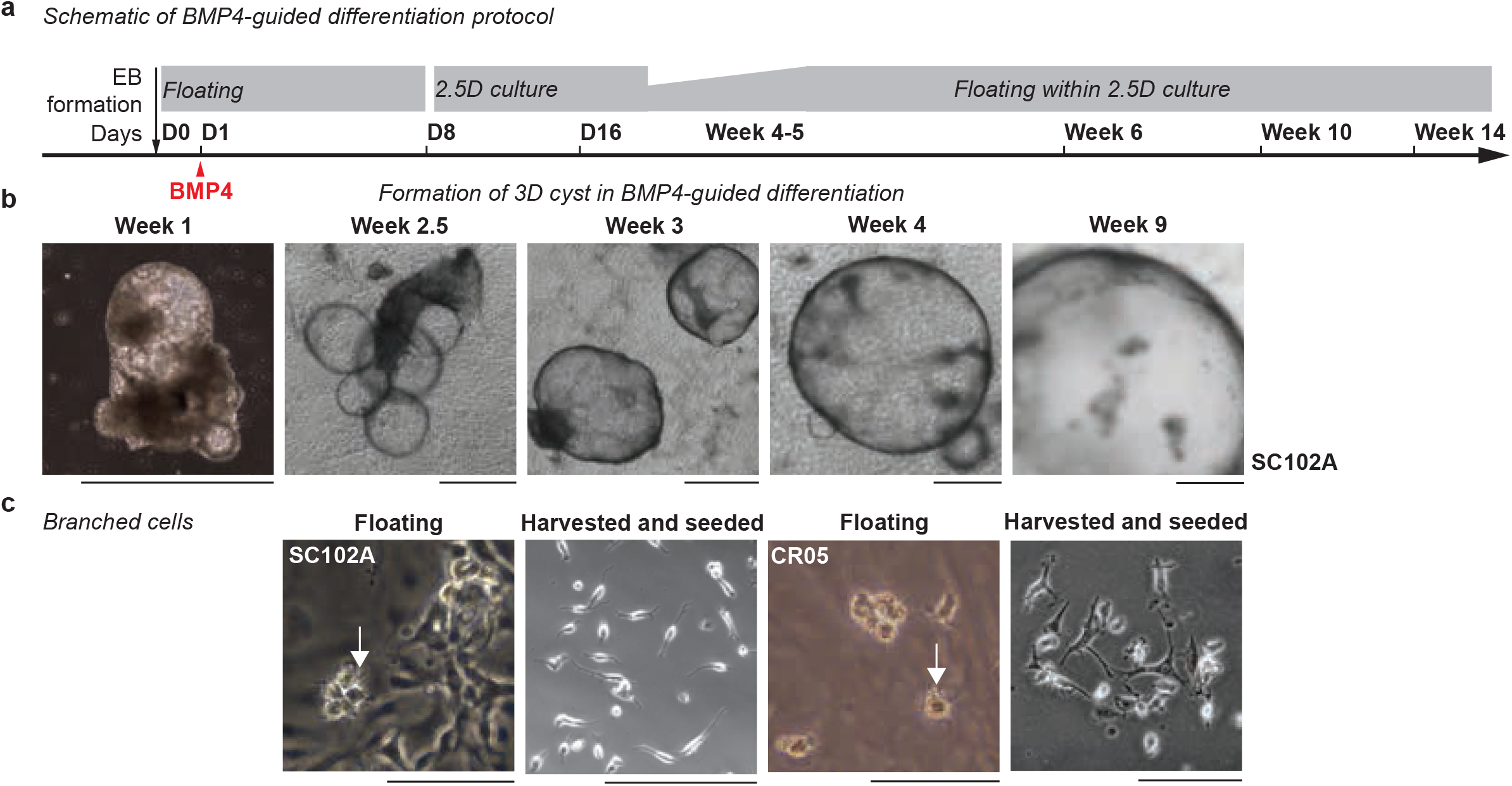
BMP4-guided differentiation induces 3D-cyst development. **a**, Schematic of differentiation protocol with a single BMP4 (bone morphogenetic protein 4) application on Day (D) 1 after induced differentiation. EB, embryoid bodies. **b**, Brightfield images of developing 3D-cysts generated with BMP4-guided differentiation for SC102A. Scale bar: 1000 μm. **c**, Brightfield images of branched cells in the supernatant for SC102A (left) and CR05 (right). Left, focus on floating cells in the original plate (week 6-7). Arrow, branching. Right, harvested supernatant and seeded on a new plate (week 5-6). Scale bar: 100 μm.

**Figure 4.**
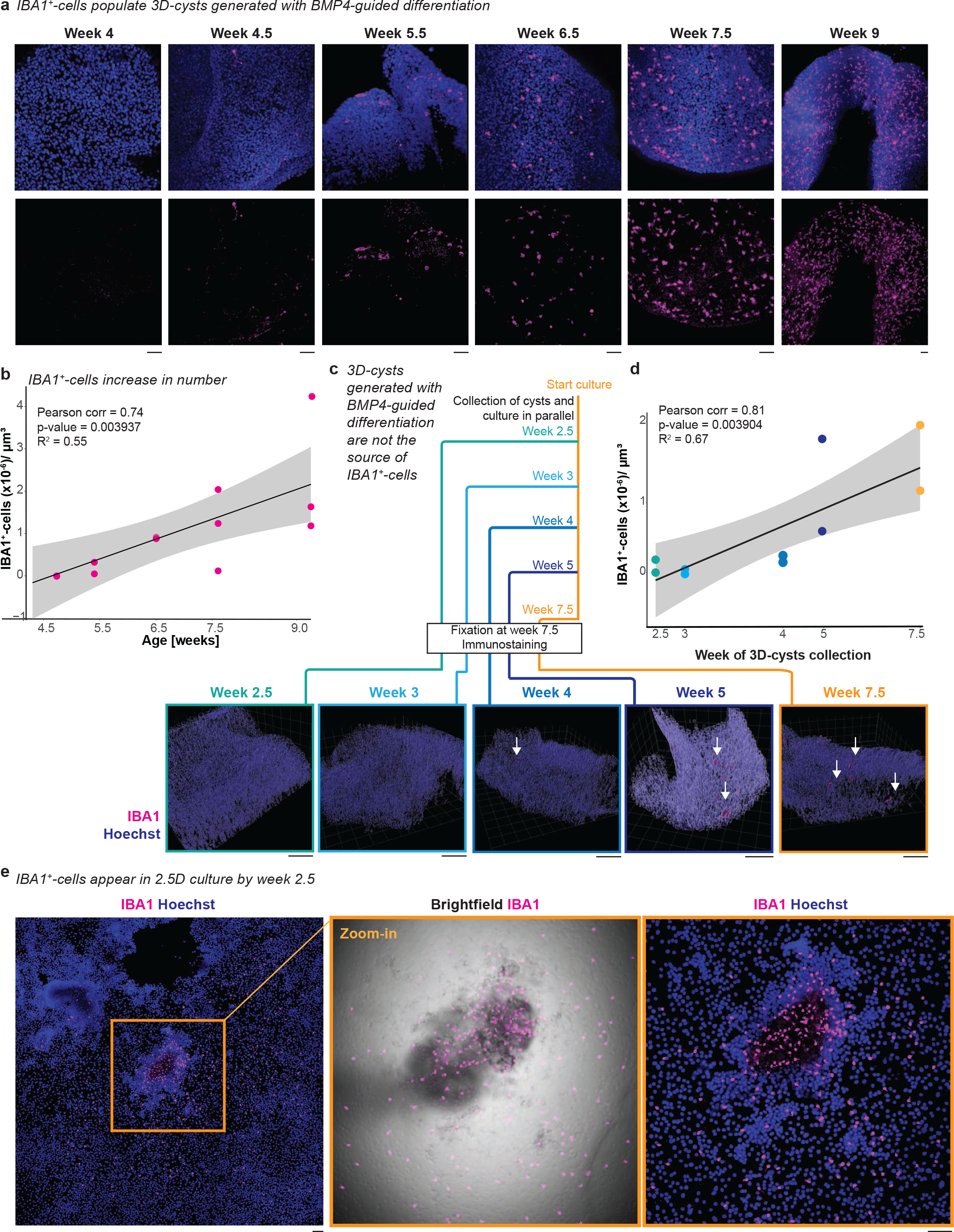
IBA1^+^-cells occupy 3D-cysts from the 2.5D plate of the BMP4-guided differentiation. Immunostaining for IBA1 (ionized calcium-binding adapter molecule 1, magenta) and Hoechst (blue) in SC102A for BMP4-guided differentiation. **a**, Timeline of the presence of IBA1^+^-cells in 3D-cysts from week 2.5 to week 9. Scale bar: 50 μm. **b**, Scatter plot of IBA1^+^-cells occupying 3D-cyst with trend curve and 95% confidence level interval. Pearson correlation showing a significant correlation between age of differentiation and number of IBA1^+^-cells occupying the cyst (Pearson’s correlation = 0.738518 and p-value = 0.003937, R^2^ = 0.5454089). **c**, 3D-cysts collected at week 2.5, 3, 4, 5 from the original plate of the 2.5D culture and whole mount immunostained together with 3D-cysts not isolated but cultured within the 2.5D culture until week 7.5 (orange frame). Representative images of 3D volume rendering to visualize the 3D-cyst with IBA1^+^-cells. White arrow, IBA1^+^-cells. Scale bar: 100 μm. **d**, Scatter plot of IBA1^+^-cells occupying isolated 3D-cyst with trend curve and 95% confidence level interval. Pearson correlation showing a significant correlation between age when 3D cysts were isolated and IBA1^+^-cells density (R = 0.81707 and p-value = 0.0039, R^2^= 0.6676). **e**, First occurrence of IBA1^+^-cells in 2.5D culture. Orange frame, zoom-in with brightfield image. Scale bar: 100 μm.

### IBA1^+^-cells associate with the mesenchymal/VIMENTIN^+^-region

Since IBA1^+^-cells similarly occupy 3D-cysts obtained with or without BMP4, we performed mass spectrometry to analyze tissue identity. Due to the reliable spatial-temporal population pattern of IBA1^+^-cells during BMP4-guided differentiation, we collected ten morphologically comparable 3D-cysts with either sparse or high IBA1^+^-cell population at week 4 and 7, respectively (**Fig. 5a**). We obtained the peptide sequence data (**Supplementary Table 2**) and compared the highly expressed proteins with 44 tissues from the human proteome atlas that have previously been characterized (Uhlén et al., 2015). At week 4, highly abundant proteins enriched across diverse tissues suggesting that cells in the 3D-cysts have various fate potentials (**Fig. 5b, Supplementary Table 3**-**4**). Interestingly, at week 7, the protein composition was specific to tissues of partial or full mesodermal origin such as soft tissue, bone marrow and smooth muscles, which is consistent with BMP4 application (**Fig. 5b, Supplementary Table 3**-**4**). Proteins specific to soft tissue were enriched, indicating the presence of cells that surround and support organs and bones such as fibrous tissue, lymph- and blood vessel, tendons, ligaments, and fat indicating a mesenchymal identity. Similarly, when we performed tissue enrichment analysis of proteins related to human eye tissue (Dunn et al., 2019; Zhang et al., 2015, 2016a), we found a significant overlap of highly abundant proteins for meninges and sclera, both of which have mesenchymal origin. In contrast, proteins enriched in ectodermal retina and optic nerve were underrepresented in our mass spectrometry data (**Fig. 5c, Supplementary Table 5**).

**Figure 5.**
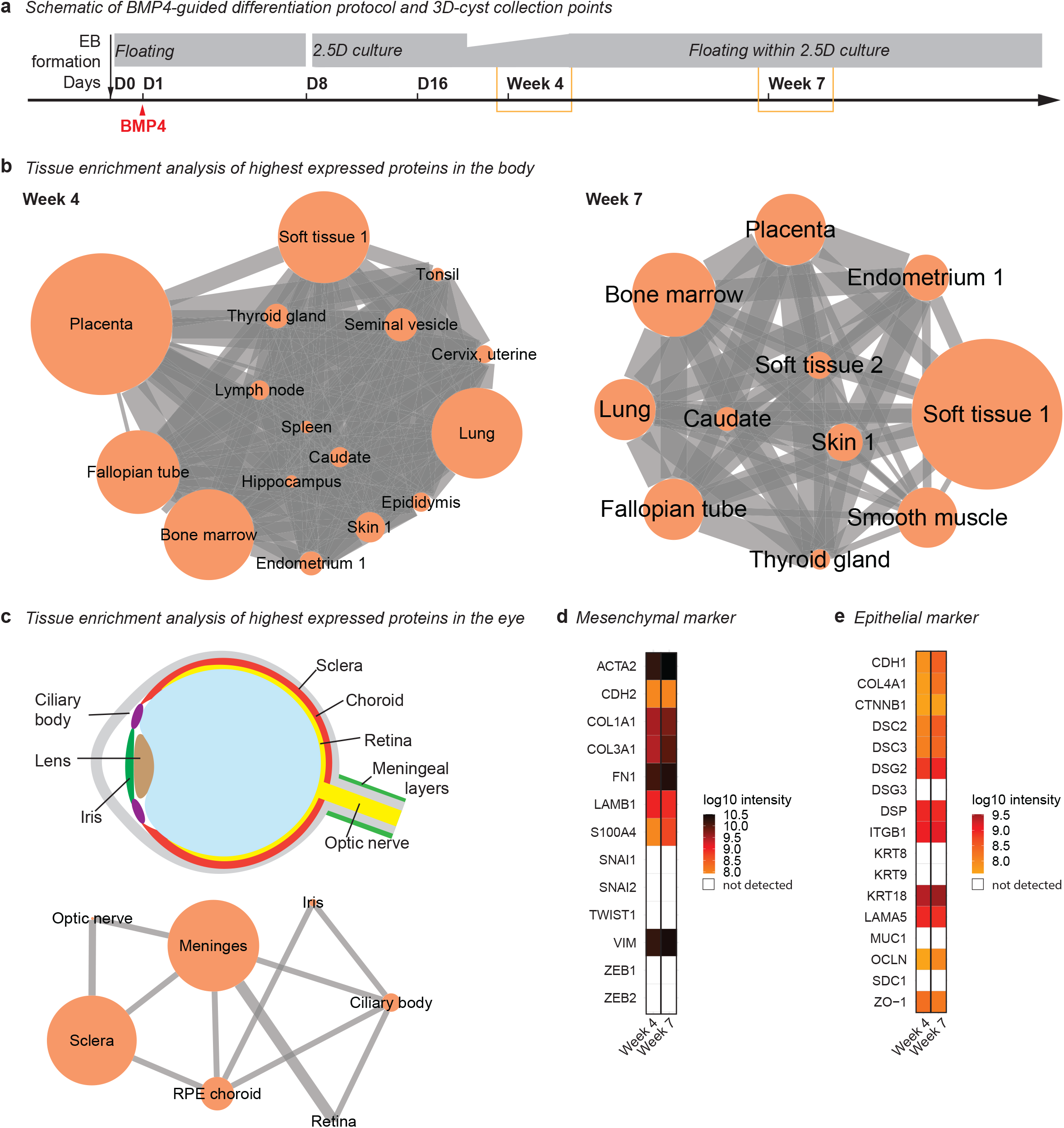
Tissue-specific protein enrichment analysis of 3D-cysts indicates a mesenchymal-like compartment. **a**, Schematic of experimental design. For mass spectrometry, ten 3D-cysts generated with BMP4-guided differentiation were collected at week 4 and week 7. **b**, Tissue enrichment analysis shown as a network. Nodes, tissue. Edges, connecting the tissues that share highly expressed proteins in the 98th percentile. The node size reflects the enrichment *p*-value and the thickness of the edge reflects the size of the shared proteins (see also: **Supplementary Table 4**). **c**, Top, eye schematic. Bottom, tissue enrichment analysis of the week 7 dataset. RPE, retinal pigment epithelium (see also: **Supplementary Table 5**). **d**-**e**, Heatmap of protein expression level in log10 (intensity) for week 4 (W4) and week 7 (W7) cystic compartments for **d**, mesenchymal marker (see also: **Supplementary Table 6**); **e**, epithelial marker (see also: **Supplementary Table 6**). White: not detected.

To validate whether the cysts are enriched for mesenchyme, we compared our mass spectrometry data with mesenchymal markers including transcription factors, cytoskeletal-, cell surface and extracellular matrix proteins (Andrzejewska et al., 2019; Owusu-Akyaw et al., 2019; Scanlon et al., 2013). We found that mesenchymal markers such as vimentin (VIM), laminin β1 (LAMB1), and fibronectin (FN1) were enriched with VIM among the most abundantly expressed proteins at week 7 (**Fig. 5d**). Since one characteristic of mesenchyme is close interaction with the epithelium (MacCord, 2012), we also investigated the presence of epithelial proteins in our dataset (**Fig. 5e**). We found several epithelial markers were expressed and upregulated from week 4 to 7. One of these epithelial markers is E-Cadherin (CDH1), which is commonly described together with the mesenchymal marker VIM in the epithelial-mesenchymal transition during development and cancer (Hay, 2005; Thiery et al., 2009; Yamashita et al., 2018). Therefore, we performed immunostaining to validate the expression of both markers in 3D-cysts derived with the BMP4-guided differentiation. VIM expression was strong in the cell layers facing the surface of the 3D-cyst wall (**Fig. 6a**). In contrast, E-Cadherin marked a defined layer next to the cystic lumen and complemented VIM expression (**Fig. 6b**). When we investigated the location of IBA1^+^-cells, they mostly occupied the VIM^+^-region (**Fig. 6c**) and stayed distinct from the E-Cadherin^+^-layer (**Fig. 6d**). Only occasionally were they observed to be intermingled with the E-Cadherin^+^-cells (**Fig. 6e**) suggesting that IBA1^+^-cells prefer the mesenchymal-like, VIM^+^-region.

**Figure 6.**
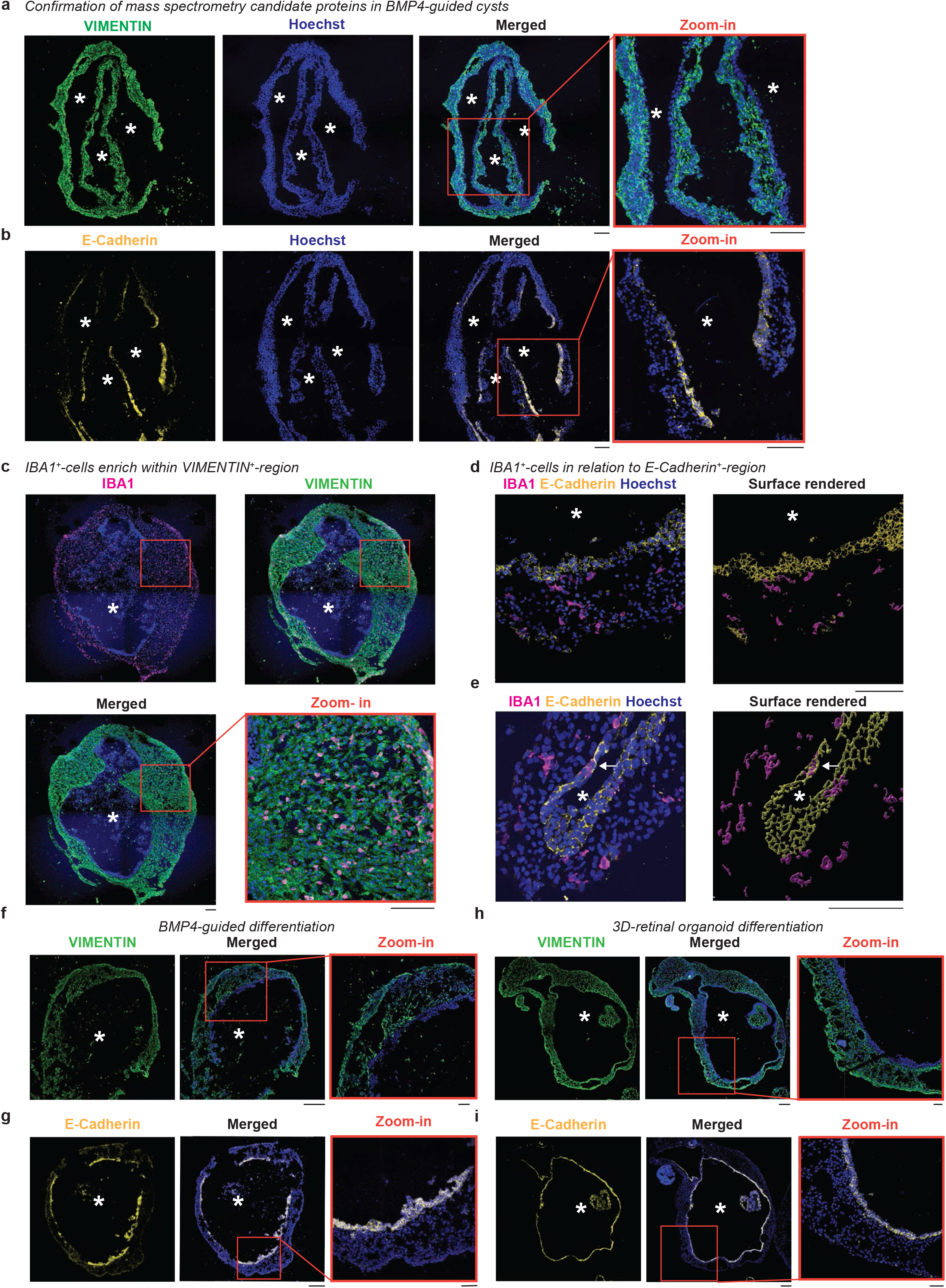
IBA1^+^-cells localize within VIM^+^-region in cystic compartment. Immunostaining of cryostat sections of cystic compartments from differentiations guided with BMP4 (**a**-**g**) or no BMP4 (**h**-**i**), counter-stained with the nuclei-dye Hoechst (blue). *, lumen. **a-e**, Scale bar: 100 μm. **a**-**b**, Sequential sections of SC102A cysts stained with VIMENTIN (green, **a**) and E-Cadherin (yellow, **b**) at week 7 with zoom-in (orange frame). **c**-**e**, Staining for IBA1 (ionized calcium-binding adapter molecule 1, magenta), VIMENTIN (green, week 12, **c**), and E-Cadherin (yellow, week 10, with 3D-surface rendering for CR05 cysts **d**-**e**). White arrow, IBA1^+^-cells within E-Cadherin layer. **f**-**i**, Comparison of immunostaining of Vimentin (green, **f, h**) and E-Cadherin (yellow, **g, i**) for SC102A at week 8 for 3D-cysts with (**f**-**g**) and without BMP4 exposure (**h**-**i**). Scale bar: 200 μm. Zoom in, Scale bar: 50 μm.

To verify whether we find a comparable pattern in the 3D-cysts obtained from the retinal organoid differentiation protocol without BMP4 (**Fig. 1a**), we repeated the above-described staining. VIM labeled a similar defined region in the 3D-cysts with E-Cadherin expression localized in a defined layer around the cystic lumen (**Fig. 6f-i**). In addition, E-Cadherin^+^-expression was found at regions facing the surface (**Fig. 7a**). IBA1^+^-cells rarely intermingled with the E-Cadherin^+^-layer facing the lumen or the surface of the 3D cyst (**Fig. 7a-b**) and mostly localized within the VIM^+^-region (**Fig. 7c**) suggesting that IBA1^+^-cells prefer the mesenchymal region.

**Figure 7.**
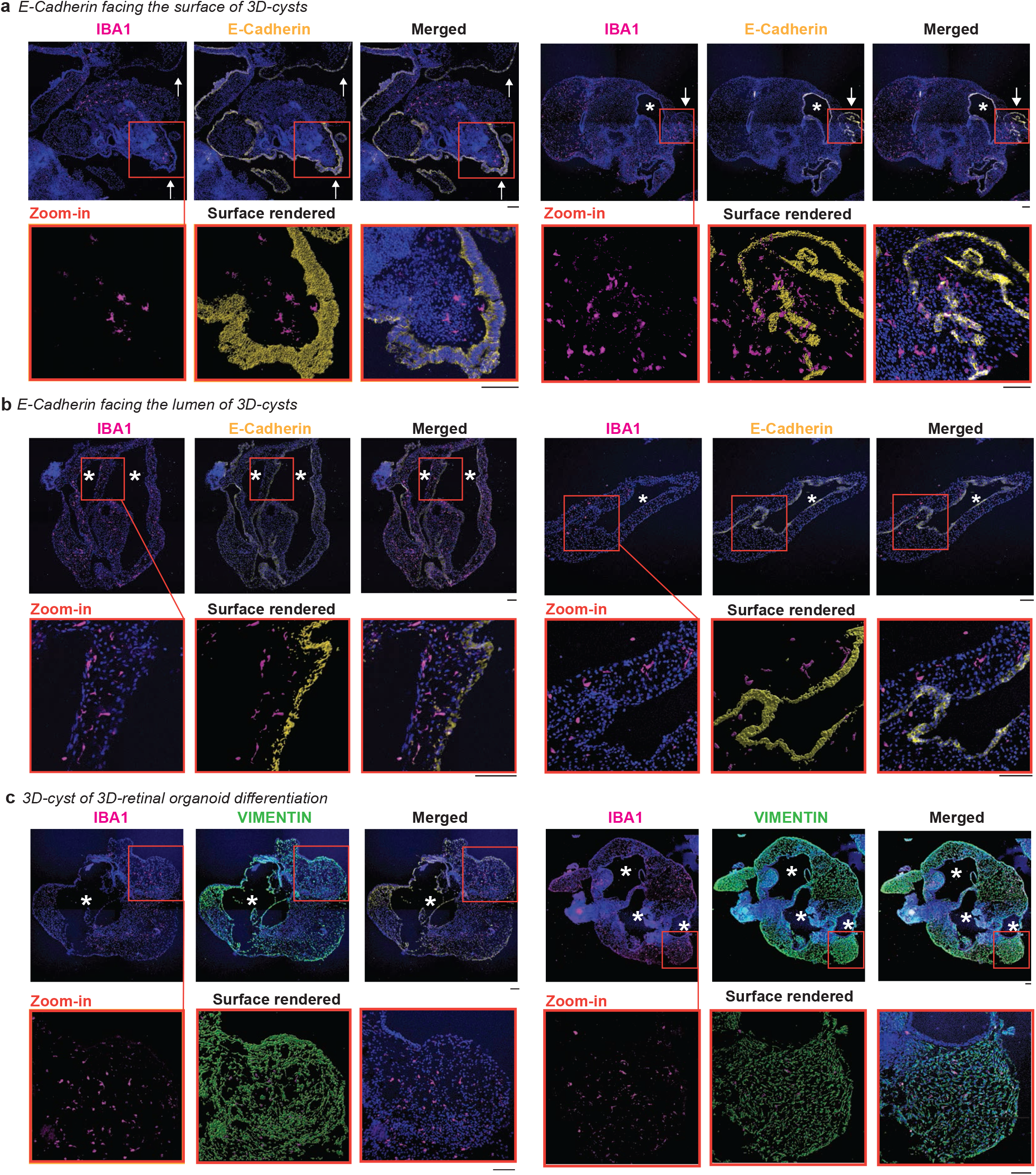
IBA1^+^-cells preferentially localize within VIM^+^-regions in 3D-cysts. Immunostaining of cryostat sections of 3D-cysts from retinal organoid differentiation without BMP4 for IBA1 (ionized calcium-binding adapter molecule 1, magenta), E-Cadherin (yellow, **a**-**b**), VIMENTIN (green, **c**), and the nuclei-dye Hoechst (blue) for SC102A at week 8. *, lumen within the cystic compartment. **a**, Arrow, E-Cadherin staining on 3D-cyst surface. Red frame, zoom-in with surface rendering in the middle. Scale bar: 100 μm.

Mesenchymal stem cells can have immunomodulatory capabilities (Andrzejewska et al., 2019) and could sequester IBA1^+^-cells away from other compartments. To test this, we first generated 2.5D cultures with the BMP4-guided differentiation containing floating IBA1^+^-cells and 3D-cysts in the supernatant and added separately generated 17-week-old 3D-retinal organoids to this culture (**Supplementary Fig. 5a**). We confirmed the development of the retinal-cup with OTX2 but we did not find IBA1^+^-cells integrated into the 3D-retinal organoid (**Supplementary Fig. 5b**). When we harvested IBA1^+^-cells from the BMP4-guided 2.5D culture, and applied them to other 3D-retinal organoids (**Supplementary Fig. 5c**), we found that IBA1^+^-cells integrated into the retinal-cup (**Supplementary Fig. 5d-e**). These data show that when there is no cystic compartment, IBA1^+^-cells start to occupy the retinal organoid. Otherwise, they are preferentially found in the mesenchymal, cystic region.

### IBA1^+^-cells adopt a BAM signature in the mesenchymal environment

The mesenchyme is involved in the development of the lymphatic and circulatory system amongst other tissues (MacCord, 2012; Pill et al., 2015; Wimmer et al., 2019). When we stained the cystic compartment for blood vessel endothelium with platelet endothelial cell adhesion molecule marker PECAM-1/CD31 (Stremmel et al., 2018; Wimmer et al., 2019), we found sparsely labeled CD31^+^-endothelial structures within the VIMENTIN^+^-region and IBA1^+^-cells intermingled within these structures (**Fig. 8a**). In contrast to mouse, human microglia infiltrate the cortex from the ventricular lumen and the leptomeninges at 4.5 gestational weeks (Monier et al., 2007; Rezaie et al., 2005), a period preceding hepatic and bone marrow hematopoiesis (Juul and Christensen, 2018; Menassa and Gomez-Nicola, 2018). Since the meninges are mesenchymal and VIM^+^ (Hardy et al., 2000), we hypothesized that IBA1^+^-cells enriched in the cysts express a signature for border-associated macrophages (BAMs). BAMs are non-parenchymal macrophages that reside either at perivascular structures, meninges, or choroid plexus. Transcriptional profiling of macrophage population in embryonic mouse brain identified CD163 as a potential marker for BAMs (Utz et al., 2020), which also labels human perivascular macrophages (Fabriek et al., 2005), mononuclear phagocytes in the choroid plexus, and cells in the meningeal- and subpial granular layer (Rezaie and Male, 2003). Indeed, we found that 99% of IBA1^+^-cells co-expressed CD163 by week 10 in the 3D-cyst (**Fig. 8b-c**). Interestingly, the onset of CD163 expression occurs in a defined window. At week 5 in the 2.5D culture, IBA1^+^-cells were still negative for CD163 (**Fig. 8d**). Within one week, IBA1^+^-cells co-expressed CD163, as they started to distribute within the 2.5D culture and occupy compartments that were sparse in nuclei (**Fig. 8e**).

**Figure 8.**
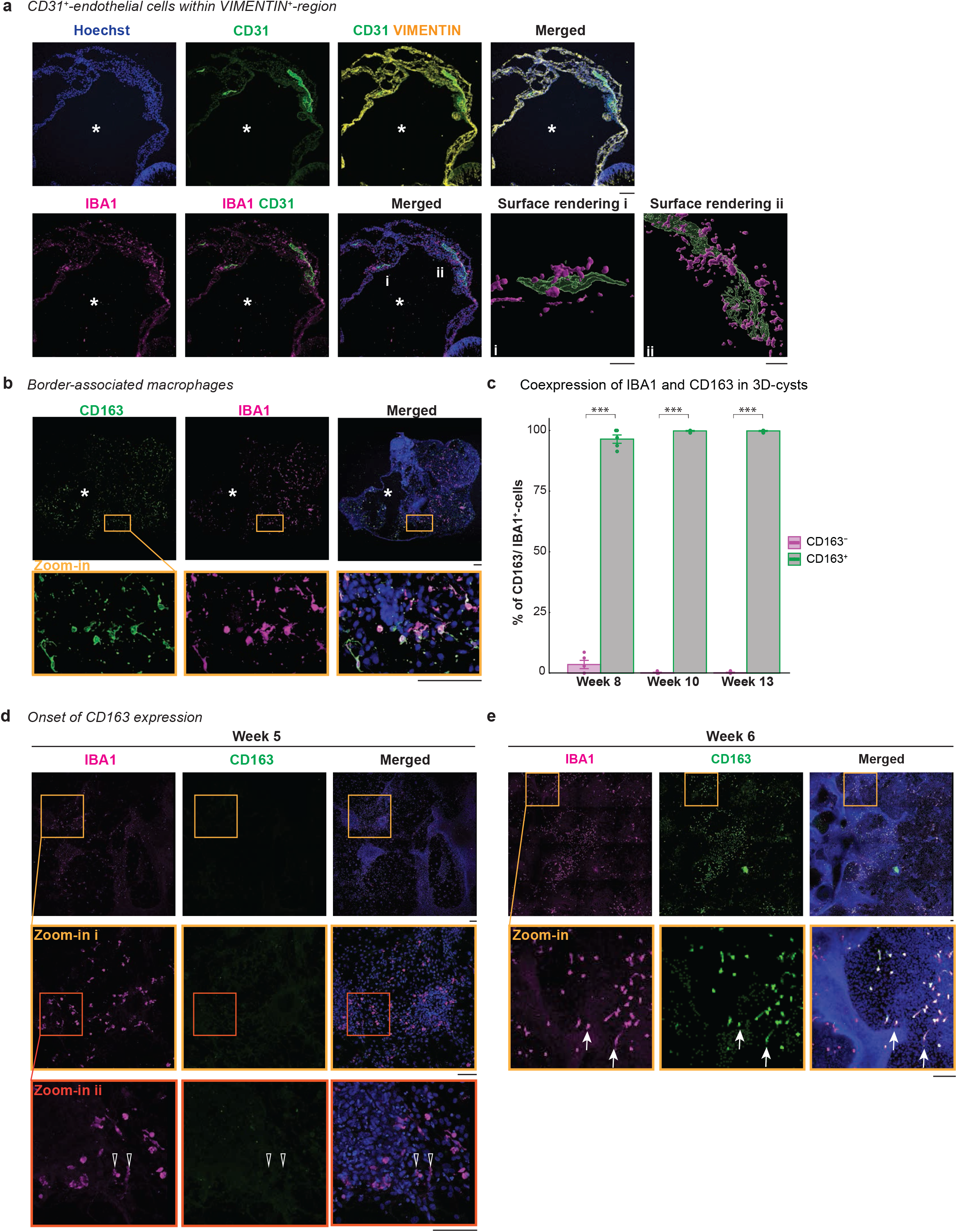
IBA1^+^-cells express CD163, a marker for border-associated macrophages. Immunostaining of cryostat sections of SC102A cysts from unguided differentiation for IBA1 (ionized calcium-binding adapter molecule 1, magenta) and the nuclei-dye Hoechst (blue). *, lumen within the cystic compartment. **a**, Immunostaining for CD31 (platelet and endothelial cell adhesion molecule 1, green) and VIMENTIN (yellow) for SC102A at week 8. Scale bar: 100 μm. Zoom-in, 3D-surface rendering (i, ii). Scale bar: 20 μm. **b**, Immunostaining of CD163 (cluster of differentiation 163/ protein tyrosine phosphatase receptor type C, green) in SC102A at week 8 with zoom-in. Scale bar: 100 μm. **c**, Bar chart of % of CD163^+^/IBA1^+^-cells and % of CD163^-^/IBA1^+^-cells per time point with standard error. Each dot represents one section of individual 3D-cyst. Two-way ANOVA p-value = < 2.2e-16 with selected post hoc-test. ^***^p < 0.001. **d**-**e**, CD163 expression in 2.5D-culture for CR05 at week 5 (**d**) and week 6 (**e**). Open arrow, CD163^-^/IBA1^+^. Arrow, CD163^+^/IBA1^+^. Red frame, zoom in. Scale bar: 100 μm.

This finding suggests that the mesenchymal environment alters the microglia signature towards BAMs, and reflects an early stage of microglia infiltration and adaptation.

## Discussion

In this study, we demonstrate that microglia-like cells emerge between week 3 to 4 in 2.5D culture (**Fig. 1c**) in an unguided retinal organoid differentiation protocol (Zhong et al., 2014). At this time point, the 2.5D culture is highly heterogenous and reflects an unperturbed intrinsic self-organized patterning environment. The properties of EBs allow the formation of retinal cell types derived from the ectodermal lineage (**Supplementary Fig. 1d**) as well as the development of mesenchymal regions mostly derived from the mesoderm (**Fig. 6h, 7**) (MacCord, 2012). This environment and the time frame is similar to when IBA1^+^-cells have been reported to appear in human embryonic development (Bloom and Bartelmez, 1940; Kelemen and Jánossa, 1980; Monier et al., 2007; Rezaie et al., 2005). Our results also support a recent study that identified a cluster of microglial cells using RNA-sequencing of brain organoids with bilateral optic vesicles (Gabriel et al., 2021). However, in contrast to human retinal development (Hu et al., 2019), we rarely observed microglia-like cells in the retinal cup at 5 weeks or later (**Supplementary Fig. 3a**). Instead, IBA1^+^-cells strongly preferred the mesenchymal-like cystic-over the neuronal compartment (**Fig. 2**).

In humans, microglia first enter the embryonic cortical regions via the ventricular lumen, choroid plexus, and leptomeninges (Menassa and Gomez-Nicola, 2018; Monier et al., 2007; Rezaie and Male, 2003), all tissues originating from the mesenchyme (Catala, 2019; Lopes, 2009; O’Rahilly and Müller, 1986). As mesenchymal structures develop *in vivo* around the neuronal retina with the choroid close to the photoreceptors and meninges wrapping the optic nerve (Forrester et al., 2010; Sturrock, 1987), the preferential location of our IBA1^+^-cells in the mesenchymal region potentially recapitulates how microglia enter the retina. The CD163 expression of IBA1^+^-cells further supports a perivascular-associated role. Initially, parenchymal microglia and border-associated macrophages are derived *in vivo* from the same primitive macrophage precursor (Goldmann et al., 2016) and then adapt their transcriptional landscape to their local environment at early developmental stages (Gosselin et al., 2017; Masuda et al., 2019; Utz et al., 2020). CD163 is one example of a human border-associated- (perivascular, leptomeningeal, choroid plexus) macrophage marker (Fabriek et al., 2005; Rezaie and Male, 2003), and its expression is upregulated during mouse embryonic development (Utz et al., 2020). Indeed, our IBA1^+^-cells in the 2.5D culture arose as CD163 negative and express CD163 after one week (**Fig. 8d-e**).

However, it remains unclear why IBA1^+^-cells do not further infiltrate the neuronal compartment especially as they are in direct contact within the 3D-retinal organoid attached cysts (**Fig. 2d**). Even if we transferred 3D-retinal organoids to 2.5D cultures of BMP4-guided differentiation, IBA1^+^-cells favor the cystic-over the neuronal compartment (**Supplementary Fig. 5**). It is possible that the cystic compartment releases guidance signals that attract IBA1^+^-cells. These cues are likely to be similar to those that recruit macrophages to the epithelial-mesenchymal transition sides in glioma (Song et al., 2017). The cystic compartment might represent a choroid plexus-like structure as we frequently observed a cuboidal epithelial surface (**Fig. 2a**), and it has been described that the choroid plexus has both an ectodermal and mesodermal compartment (Javed et al., 2021). However, our staining for choroid plexus structures with transthyretin were not reliable (data not shown), which we suspect is due to the rapidly changing environment. There is also limited knowledge about developmental guidance cues that attract microglia to the neuronal compartment and more specifically to the retina. Further investigation will be required to answer these questions.

A possible limitation of our model is the intrinsic variability of the unguided differentiation that currently prevents production of a high number of cystic compartments and microglia-like cells, which likely depends on BMP4. Further optimization will allow the advantage of unguided differentiation while increasing the likelihood of microglia occurrence. Another limiting factor are the hiPSC lines used for differentiation. In our study, we employed two hiPSC lines from different origins and both resulted in a similar phenotype. We cannot exclude that the qualitative outcome could be different for other hiPSC lines due to intrinsic properties related to genetic origin, epigenetic landscape, or transcriptional state at the time of the differentiation (Kilpinen et al., 2017; Ortmann and Vallier, 2017). Such factors might prevent the generation of cystic compartments and therefore the appearance of microglia-like cells.

In summary, our study confirms that microglia-like cells occur during retinal organoid differentiation and preferentially occupy mesenchymal region. These findings will allow future analysis of microglial migration in complex tissue environments and facilitate identification of mechanistic cues that attract microglia in complex tissue structures.

## Figure legends

**Supplementary Table 1 - Overview of human induced pluripotent stem lines included in this study.**

hPSCreg.eu, human pluripotent stem cell registry. MYC, MYC Proto-Oncogene. KLF4, Kruppel Like Factor 4. Large T antigen, large tumor antigen. LIN28, Zinc Finger CCHC Domain-Containing Protein. OCT4 (Octamer-Binding Protein 4)/ POU5F1 (POU Domain, Class 5, Transcription Factor 1). SOX2, Sex Determining Region Y-Box 2. SV40, Simian-Virus 40.

**Supplementary Table 2 – Mass spectrometry raw data.**

**Supplementary Table 3 – Mass spectrometry for tissue enrichment for Fig. 5b-c.**

**Supplementary Table 4 – Tissue enrichment analysis enhanced tissues for Fig. 5b.**

**Supplementary Table 5 – Eye tissue enrichment analysis for Fig. 5c.**

**Supplementary Table 6 – Mesenchymal and epithelial marker for Fig. 5d-e.**

**Supplementary Table 7 – Antibody list.**

## Author contributions

Conceptualization: S.S.; Data curation and Formal analysis: V.H., R.J.A.C. Methodology: K.B., V.H., M.K., R.J.A.C., S.S.; Validation and Investigation: K.B., V.H., M.K.; Resources: A.V., K.R., T.C.; Writing – Original draft and Visualization: V.H., S.S. with input from K.B., M.K.; Supervision and Funding Acquisition: S.S.

Acknowledgements

We thank the scientific service units at IST Austria specifically the life science facility and bioimaging facility for their support; Nicolas Armel for performing the Mass Spectrometry. We thank Alexandra Lang and Tanja Peilnsteiner for their help in human brain tissue collection. We thank all members of the Siegert group for constant feedback on the project and Margaret Maes, Rouven Schulz, and Marco Benevento for feedback on the manuscript. This research was supported by the Gesellschaft für Forschungsförderung Niederösterreich (grant No. Sc19-017 to V.H.) and the European Research Council (grant No. 715571 to S.S.).

## Methods

### Ethical approval

The IST Austria Ethics Officer and Ethics Committee approved the use of human induced pluripotent stem cells (hiPSC). The use of human brain samples was approved by the Ethics Committee of the Medical University Vienna.

### Primary human tissue samples

Human brain samples were explanted from the temporal cortex (T1) of patients undergoing temporal lobe surgery for epilepsy treatment. Immediately after the surgical explant, the samples were transferred into saline solution (0.9% (v/v) NaCl (Braun 3570160) in H_2_O). The tissue was immersed in 4% (w/v) PFA within 5 minutes and post-fixed on an orbital shaker at 4ºC overnight.

### Cell lines

This study used two human induced pluripotent stem cell lines: SC 102A-1 GVO-SBI Human Fibroblast-derived (feeder-free) iPSC cell line (BioCat; male; hPSCreg.eu: SBLi006-A; in this study referred to SC102A). NCRM-5 (aka NL-5; human umbilical cord blood CD34^+^ cells derived; RUCDR Infinite Biologicals, Cell line ID: CR0000005, NHCDR ID: ND5003; male; hPSCreg.eu: CRMi001-A; in this study referred to CR05). For more details, see (**Supplementary Table 1**).

### Cell culture and retinal organoid generation

#### Matrigel-coating

Matrigel (Corning® Matrigel® hESC-Qualified Matrix, *LDEV-Free, (Corning, #354277) was used according to the manufacturer instructions with the following modifications: Matrigel aliquots were dissolved in ice-cold X-Vivo 10 chemically defined, serum-free hematopoietic cell medium (Lonza, #BE04-380Q) prior coating the plates. 6-cm dishes (VWR, #734-0007) were coated for retinal organoid or microglia-like differentiation and 2-well chambered coverslips (Ibidi, #80286) for 2.5D culture.

#### Maintenance of human induced pluripotent stem cells

hiPSCs were cultured at 37ºC and 5% CO_2_ in a humidified incubator (BINDER C150) in mTeSR1 medium (STEMCELL Technologies, #85850) on Matrigel (Corning, #354277) coated 6-well plates (Corning, #3516). Cells were passaged in small aggregates every 3-4 days and were dissociated before reaching 80% confluency using EDTA dissociation buffer (0.5M EDTA (ethylenediaminetetraacetic acid, K.D. Biomedical, #RGF 3130), 0.9 g (w/v) NaCl (Sigma, #5886) in PBS (phosphate buffered saline, calcium/magnesium-free, Invitrogen, #14190), sterile filtered, stored at 4ºC) according to (Chen, 2014). Cells were tested on regular basis for mycoplasma using MycoAlert Mycoplasma Detection Kit (Lonza, #LT07-518). For iPSC differentiation, two wells of a 6-well plate were used for SC102A and four wells for CR05 as starting material.

#### Retinal organoid differentiation

Retinal organoids were generated as described before (Zhong et al., 2014) with the following modifications: On day 0 of differentiation, iPSC colonies were dissociated into evenly sized aggregates using a cell-passaging tool (Thermo Fisher Scientific, #23181-010). After mechanical scraping, floating aggregates were transferred with a 1250μl wide orifice pipette (VWR, #613-0737) onto one 10 cm Petri dish (Sarstedt, #82.1473), and cultured in mTeSR1 medium supplemented with 10 μM blebbistatin (Sigma, #B0560-5MG). On day 1, 2 and 3, the medium was gradually replaced with ¼, ½, and 1, respectively, of NIM (neural induction medium: DMEM/F12 (Gibco, #31331-028), 1x N2 supplement (Gibco, #17502-48), 1% (v/v) NEAA Solution (Sigma, #M7145), 2 μg/ml heparin (Sigma, #H3149-50KU). From day 4 onwards, 10 ml medium was exchanged daily with NIM. On day 8, the floating embryoid bodies (EB) were collected, equally distributed onto 8 Matrigel-coated 6-cm dishes (approximately 20-40 number of EBs/cm^2^) and cultured in 3 mL NIM. From day 16 onwards, NIM was exchanged daily for 3:1-DMEM/F12-medium (3 parts DMEM (Thermo Fisher Scientific, #31966047) and one-part F12 medium (Ham’s F-12 Nutrient Mix, Thermo Fisher Scientific, #31765-027), supplemented with 2% (v/v) B27 without vitamin A (Thermo Fisher Scientific, #121587-10), 1% (v/v) NEAA solution (Sigma, #M7145), 1% (v/v) penicillin-streptomycin (Thermo Fisher Scientific, #15140122). On day 28-32, optic-cup structures were manually micro-dissected from the 6-cm plate and transferred into a 3.5-cm Petri dish (Corning, #351008) containing 2.5 mL 3:1-DMEM/F12-medium. 3:1-DMEM/F12-medium was exchanged twice per week. From day 42 onwards, 3:1-DMEM/F12-medium was supplemented with 10% (v/v) heat-inactivated FBS (Thermo Fisher Scientific, #10270-106) and 100 μM taurine (Sigma, #T0625-25G). At week 10, the 3:1-DMEM/F12-medium was supplemented with 10 μM retinoic acid (Sigma, #R2625), and the medium was daily exchanged. At week 14, B27 supplement in the 3:1-DMEM/F12-medium was replaced with 1x N2 supplement (Gibco, #17502-48), 10% (v/v) heat-inactivated FBS (Thermo Fisher Scientific, #10270-106), 100 μM taurine (Sigma, #T0625-25G) and the retinoic acid concentration was reduced to 5 μM.

#### Retinal organoid differentiation – Maintenance beyond day 28-32 in 2.5D culture

The differentiation protocol is identical to the “retinal organoid differentiation” section with the following modifications: On day 8, EBs within a volume of 1.5 mL were transferred on Matrigel-coated 2-well chambered coverslip. After the change to the 3:1-DMEM/F12-medium on day 16, 2.5D cultures were exclusively maintained in this media with daily media changes without any additional supplements that are typically added at later differentiation time points in the “retinal organoid differentiation”.

#### BMP4-guided cystic compartment and microglial-like cell differentiation

The differentiation protocol is identical to the “Retinal organoid differentiation – Maintenance beyond day 28-32 in 2.5D culture” section with the following differences: On day 1, 12.5 ng/mL (final concentration) of recombinant human BMP4 (Peprotech, #120-05) was added as a single shot. From D8 onwards, medium was exchanged twice per week.

#### Harvesting microglia-like cells after BMP4 application

From D40 onwards, microglia-like cells released into the supernatant were harvested. For this, the supernatant was collected and centrifuged (VWR, Mega Star 3.0R) at 200g for 4 minutes. Cells were resuspended in 3:1-DMEM/F12-medium, and transferred into 8-well chambers (IBIDI, #80826) for immunostaining.

#### Harvesting cystic structure after BMP4 application

At D18, D21, D28, D35 floating cystic structures were transferred into a new 3.5 cm petri dish (Corning, #351008) using a 1250μL wide orifice pipette tip and cultured in 2 mL 3:1-DMEM/F12-medium in parallel to not-transferred cysts, which were further cultured in the original differentiation dish until D45. The medium was exchanged twice per week until D45 when all time points were fixed as described in the result section.

#### Culturing retinal organoids within BMP4-guided cystic compartments

At D118, eight retinal organoids were transferred into a dish containing BMP4-guided cystic compartment and microglia-like cells and cultured for 10 days. The medium was exchanged to 3:1-DMEM/F12-medium supplemented with 1x N2 Supplement, 10% (v/v) heat-inactivated FBS, and 100 μM taurine (Sigma, #T0625-25G). 3 mL medium was exchanged twice per week and 5 μM retinoic acid was added daily.

#### Supplementing retinal organoids with microglia-like cells

At D118, eight retinal organoids were transferred into a 24 well plate. Microglia-like cells were harvested as described “Harvesting microglia-like cells after BMP4 application” from two 6 cm dishes and added to the organoids once. The medium was exchanged to 3:1-DMEM/F12-medium supplemented with 1x N2 Supplement, 10% (v/v) heat-inactivated FBS, and 100 μM taurine (Sigma, #T0625-25G). Organoids and microglia-like cells were cultured for 10 days. 2 mL medium was exchanged twice per week and 5 μM retinoic acid was added daily.

### Histology

#### Histology - human brain samples

After PFA fixation, the samples were washed with PBS at least for 15 minutes three times. The samples were embedded in 3% (w/v) agarose (Sigma, #A9539) and sliced with a vibratome (Leica VT 1200) at a thickness of 100 μm. The vibratome slices were then cryoprotected with 30% (w/v) sucrose (Sigma, #84097, sterile filtered) until they sunk in the solution. The samples were stored at -80ºC until further use.

#### Fixation of retinal organoids/cystic structures (=aggregates)

Aggregates were fixed in 4% (w/v) PFA in PBS for 20 minutes at room temperature, then washed three times with PBS at room temperature and cryopreserved in 30% (w/v) sucrose in PBS overnight at 4 ºC.

#### Cryostat sectioning

Cryopreserved aggregates were transferred to a cryomold (PolyScience, #18985) using a 1250μL wide orifice pipette tip and embedded in Tissue-Tek O.C.T. compound (TTEK, A. Hartenstein) on dry ice. Samples were stored at -80ºC until further use. Cryosections (20-30 μm) of aggregates were generated using a cryostat (MICROM, NX70 CRYOSTAR, Thermo Scientific). Sections were mounted onto Superfrost Plus glass slides (Lactan, #H867.1), dried at room temperature overnight and stored at -80ºC until further use. For immunostaining, slides were dried for 1 h at room temperature. Sections on glass slices were encircled with an engraving, hydrophobic pen (Sigma-Aldrich, #Z225568).

#### Immunostaining of cryostat sections

Cryostat sections were incubated in “blocking solution” containing 1% (w/v) bovine serum albumin (Sigma, #A9418), 5% (v/v) Triton X-100 (Sigma, #T8787), 0.5% (w/v) sodium azide (VWR, #786-299), and 10% (v/v) serum (either goat, Millipore, #S26, or donkey, Millipore, #S30) for two hours in a humidified chamber protected from light at room temperature. Afterwards, the samples were immunostained with primary antibodies diluted in antibody solution containing 1% (w/v) bovine serum albumin, 5% (v/v) triton X-100, 0.5% (v/v) sodium azide, 3% (v/v) goat or donkey serum, and incubated overnight in a humidified chamber at room temperature. For the list of primary antibodies: **Supplementary Table 7**.

The sections were washed three times with PBS and incubated in a light-protected humidified chamber for 2 hours at room temperature, with the secondary antibodies diluted in antibody solution. The secondary antibodies raised in goat or donkey were purchased from Thermo Fisher Scientific (Alexa Fluor 488, Alexa Fluor 568, Alexa Fluor 647, 1:2000). The sections were washed three times with PBS. The nuclei were labeled with Hoechst 33342 (Thermo Fisher Scientific, Cat#H3570, 1:5000 diluted in PBS) for 8 minutes, and after a final two times PBS wash embedded using an antifade solution [10% (v/v) mowiol (Sigma, #81381), 26% (v/v) glycerol (Sigma, #G7757), 0.2M tris buffer pH 8, 2.5% (w/v) Dabco (Sigma, #D27802)] with microscope cover glass slips (Menzel-Glaser #0). Samples were stored at 4 ºC until imaging.

#### Immunostaining for human brain slices

The staining was performed as described under “Immunostaining for cryostat sections” with following adaptations: Floating brain slices were stained in a 24-well plate and the primary antibody was incubated for 48 hours on a shaker. After immunostaining, the slices were mounted on glass microscope slides (Assistant, #42406020) and embedded with antifade solution.

#### Immunostaining for whole mount aggregates

The staining was performed as described under “Immunostaining for cryostat sections” with the following adaptations: The primary antibody was incubated for 48 hours on a shaker at room temperature, and washed at least for two hours.

#### Immunostaining for whole mount organoids

The staining was performed as described under “Immunostaining for cryostat sections” with the following adaptations: Organoids were incubated in blocking solution for 2 days on a shaker at 4 ºC. The primary antibody concentration was doubled and organoids were incubated for 10 days on a shaker at 4 ºC, and washed three times in PBS at least for one day. Then the organoids were incubated with secondary antibodies (1:500) and Hoechst (1:1000) diluted in antibody solution simultaneously for 3 days on a shaker at 4ºC. After washing the organoids three times in PBS for one day, organoids were mounted with low gelling agarose followed by a glycerol gradient as described in “Immunostaining for whole mount aggregates”.

#### Mounting of whole mount aggregates

For whole mount aggregates, the tissue was mounted on 8-well chambers (IBIDI, #80826) using 3% (w/v) low gelling temperature agarose (Sigma-Aldrich, #A9414-25G). Then a glycerol gradient was performed starting with 50% (v/v) glycerol (Sigma-Aldrich, G7757-1L) in H_2_O followed by 75% (v/v) glycerol in H_2_O. Afterwards, the whole mount aggregate was imaged.

### Imaging

*Brightfield.* Differentiation was monitored with a bright-field microscope (Olympus CKX41) with 5x, 10x and 20x objectives (Olympus) and a lens marker (Nikon), and an EVOS microscope (Thermo Fisher Scientific) with 2x, 4x, 10x, 20x, 40x objectives (Thermo Fisher Scientific).

#### Confocal microscopy

Images were acquired with a Zeiss LSM880 Airyscan upright or inverted or with a Zeiss LSM800 upright. Ibidi plates were exclusively imaged using an inverted microscope. For overview images Plan-Apochromat 10x air objective NA 0.45 (WD=2.1mm) or Plan-Apochromat 20x Air objective NA 0.8 were used and tile-scan z-stacks were acquired. For detailed images Plan-Apochromat 40x oil immersion objective NA 1.3 was used.

#### Image analysis

Confocal images were converted to .ims files using the Imaris converter and imported to Imaris 9.3 (Bitplane Imaris 3/4D Image Visualization and Analysis Software).

#### Surface rendering

were generated using the surface rendering module with the surface detail set to 0.2 μm.

#### Determining the volume of organoids

The Hoechst channel was processed using the normalize layer function of Imaris. Then, a surface rendering was performed and the total volume of the Hoechst channel was determined.

#### Determining the number of IBA1^+^-cells

The spot function of Imaris was used to analyze the number of IBA1^+^-cells. The estimated XY diameter was set to 15 μm.

#### Graphics

All graphics were generated using R (version 4.1.0). Excel files were loaded into R via the xlsx package (version 0.6.1) (Dragulescu, 2014). Plots were made using ggplot2 (version 3.0.0) (Wickham, 2016). Linear regression was performed using the lme4 package (version 1.1-17) (Bates et al., 2015).

### Mass spectrometry

10 cystic structures were harvested at D28 and D45, washed once in DPBS (Thermo Fisher Scientific, # 14190-250) and snap frozen in liquid nitrogen. Samples were stored at -80°C until further analysis. For Liquid chromatography - mass spectrometry (LCMS) analysis, pelleted cystic structures were denatured, reduced, alkylated with iodoacetamide and trypsin-digested into peptides using a commercial in-Stage Tips kit (P.O.00001, Preomics), following exactly the manufacturer’s instructions. Cleaned-up, reconstituted peptides were then analyzed by Liquid chromatography – tandem mass spectrometry (LC-MS/MS) on an Ultimate High-performance liquid chromatography (HPLC) (ThermoFisher Scientific) coupled to a Q-Exactive HF (ThermoFisher Scientific). Each sample was concentrated over an Acclaim PepMap C18 pre-column (5 μm particle size, 0.3 mm ID x 5 mm length, ThermoFisher Scientific) then bound to a 50 cm EasySpray C18 analytical column (2 μm particle size, 75 μm ID x 500 mm length, ThermoFisher Scientific) and eluted over the following 180 min gradient: solvent A, water + 0.1% formic acid; solvent B, 80% acetonitrile in water + 0.08% formic acid; constant 300 nL/min flow; B percentage: start, 2%; 155 min, 31%; 180 min, 44%. Mass spectra were acquired in positive mode with a Data Dependent Acquisition method: FWHM 20s, lock mass 445.12003 m/z; MS1: profile mode, 120,000 resolving power, AGC target 3e6, 50 ms maximum IT, 380 to 1,500 m/z; MS2: top 20, centroid mode, 1.4 m/z isolation window (no offset), 1 microscan, 15,000 resolving power, AGC target 1e5 (minimum 1e3), 20 ms maximum IT, 200 to 2,000 m/z scan range, NCE 28, excluding charges 1 and 8 or higher, 60s dynamic exclusion.

Raw files were searched in MaxQuant 1.6.5.0 against the reference *Homo sapiens* proteome downloaded from UniProtKB. Fixed cysteine modification was set to Carbamidomethyl. Variable modifications were Oxidation (M), Acetyl (Protein N-term), Deamidation (NQ), Gln->pyro-Glu and Phospho (STY). Match between runs, dependent peptides and second peptides were active. All FDRs were set to 1%.

#### Tissue enrichment analysis

To determine human tissues that resemble the highly expressed protein profile in the cystic structure, we performed a tissue enrichment analysis similar to previous approaches (Angeles-Albores et al., 2016; Jain and Tuteja, 2019). Proteins specific to a given tissue (or *tissue-specific proteins*) were downloaded from the Human Protein Atlas (HPA, http://www.proteinatlas.org) (Uhlén et al., 2015) which has curated the expression profiles of human genes both on the mRNA and protein level in 44 normal human tissue types (corresponding to 62 tissue samples). In particular, tissue-specific proteins were defined as proteins that were highly detected, i.e., a strong immunohistochemical staining intensity in 25-75% of cells as annotated in HPA. Cystic-specific proteins in either D28 or D45, on the other hand, were obtained by taking proteins whose expressions are within the 98^th^ percentile of the protein expression distribution in the mass spectroscopy data. The overlap between the 3D-cyst specific proteins and the tissue-specific proteins were calculated and the hypergeometric test was used to calculate the enrichment of this overlap as

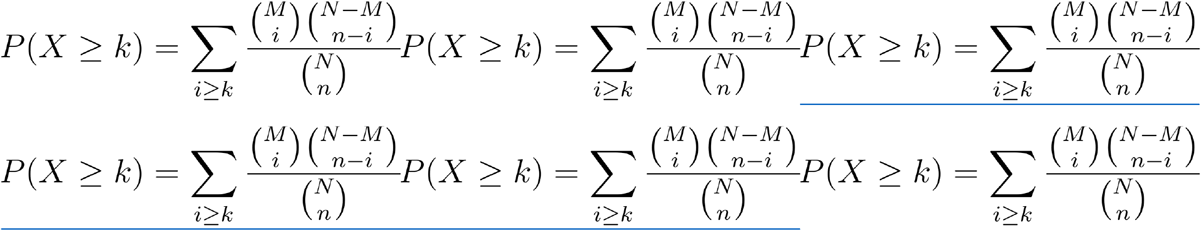

with *n* as the number of 3D-cyst specific proteins from the *N* total number of detected proteins with mass spectrometry, *M* as the number of tissue-specific proteins and *k* is the size of their overlap. The obtained *p*-values were Bonferroni-corrected for multiple comparisons implemented through the multitest function of statsmodels (Seabold and Perktold, 2010). A similar analysis was conducted for proteins found in previous mass spectrometry analyses of human iris, ciliary body, RPE/choroid (Zhang et al., 2016a), optic nerve, sclera (Zhang et al., 2016b), retina (Zhang et al., 2015) and meninges (Dunn et al., 2019). Tissue-specific protein profiles were defined as the proteins that are present in the 80th percentile of the protein expression distribution. Tissue-specific proteins that are in at most two tissues were discarded to account for possible non-specific expression. Note that while the tissue enrichment p-values change with the percentile cut-off, the qualitative results remain the same.

#### Heatmap of mesenchymal stem cell markers

The list of markers of epithelial and mesenchymal markers was obtained from (Andrzejewska et al., 2019; Owusu-Akyaw et al., 2019; Scanlon et al., 2013). Protein expression in 3D cysts for these markers was plotted as a heatmap for week 4 and 7. Fold-change was determined by dividing the intensity at D45 with the intensity at D28. Upregulated proteins are those with fold-change greater than or equal to 2.0 while downregulated proteins are those with fold-change less than or equal to 0.5.

### Statistical analysis

All statistical tests were performed using R. Models were generated by changing the default contrast for unordered variables to “contr.sum” to apply type III ANOVA to the model to evaluate the overall contribution of the response variable. Post-hoc tests were performed via the “dplyr” package (version 1.0.7) (Wickham et al., 2021) and the “multcomp” package and were corrected for multiple testing using the single-step method (Hothorn et al., 2008). Pearson correlation was performed using the “ggpubr” package (version 0.4.0) (Kassambara, 2017).

#### 3D cyst occupation and isolation

We performed a Pearson correlation test to the correlation between IBA1+-cell density and age of differentiation (**Fig. 4b, d**).

#### Co-expression of IBA and CD163

We performed two-way ANOVA to examine changes in expression over time by using an interaction of these two predictors. A random effect (cyst ID) was included to account for the dependency of the data which results from repeated counting of the same sections. As a significant effect (p < 0.05) was observed for the interaction we performed a post-hoc analysis for pair-wise comparison using the Tukey-Test and p-values were adjusted using the method set to “BH” (**Fig. 8c**).

#### Integration of IBA1^+^-cells into 3D retinal organoids

A Shapiro-Wilk test determined that the data was not normally distributed. Therefore, we performed a Wilcoxon-test to test differences between experimental conditions (**Supplementary Fig. 5d**).

#### Repetition

All experiments were performed by at least two experimentalists independently for both cell lines with exceptions of the mass spectrometry (**Fig. 5**) and (**Fig 4a-d**). In total, we performed for the hiPSC lines SC102A 18x and CR05 10x retinal organoid differentiations and for SC102A 10x and CR05 8x BMP4-guided differentiation.

---

**Supplementary Figure 1.**
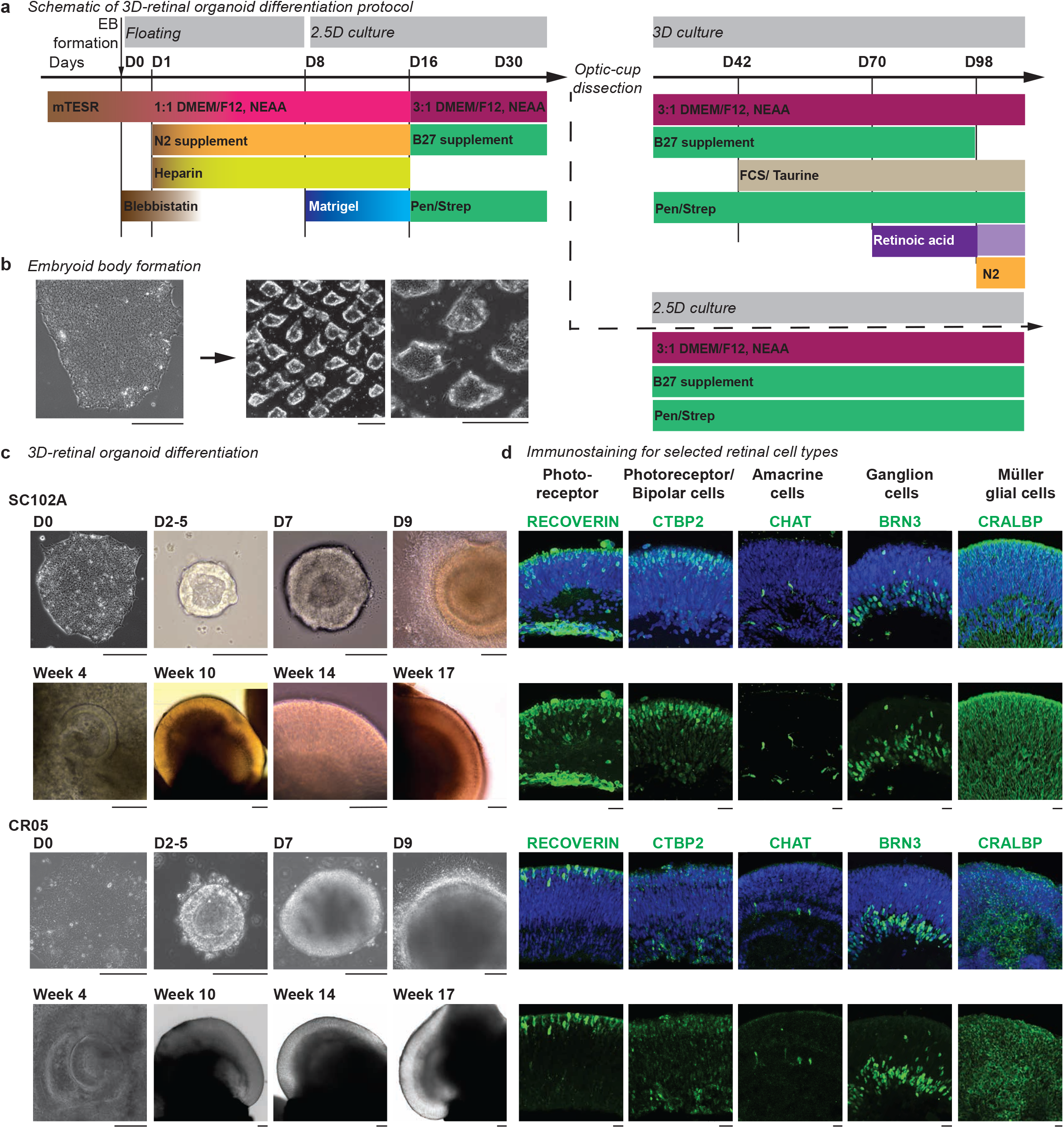
Differentiation of hiPSC lines SC102A and CR05 into 3D-retinal organoids. **a**-**b**, Schematic of retinal organoid differentiation protocol for 2.5D and 3D culture. After reaching 80% confluency, human induced pluripotent stem cells (hiPSC) were cut into evenly sized aggregates to form embryoid bodies (EBs) (**b**, Scale bar: 100 μm). EBs were cultured in suspension and were seeded on D8 on Matrigel-coated plates. At D30-32, optic-cup structures were manually micro-dissected and cultured in suspension. D, days after induced differentiation. FCS, fetal calf serum. NEAA, Non-Essential Amino Acid. Pen/Strep, penicillin and streptomycin. **c**, Brightfield images at selected days and weeks after induced differentiation (D) for SC102A (top) and CR05 (bottom). Scale bar: 100 μm. **d**, Immunostaining of cryostat sections for selected retinal cell type marker (green) and the nuclear dye Hoechst (blue) with focus on the retinal-cup for SC102A (top) and CR05 (bottom) between week 18-20 with exception of BRN3 for SC102A at week 9. CTBP2, C-Terminal-Binding Protein 2. CHAT, choline acetyltransferase. BRN3, brain-specific Homeobox/POU Domain Protein 3. CRALBP, cellular retinaldehyde-binding protein. Scale bar: 20 μm.

**Supplementary Figure 2.**
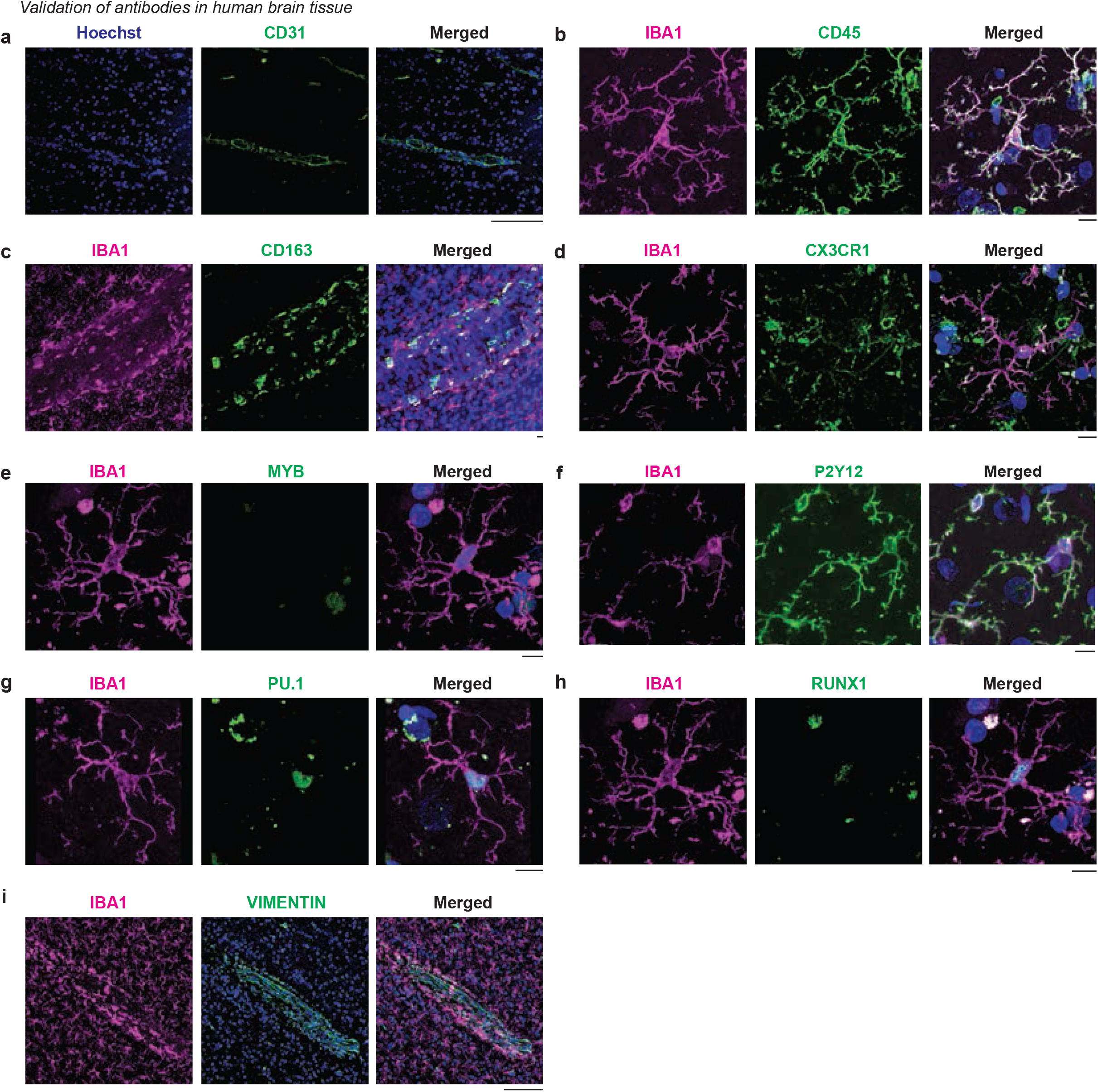
Confirmation of antibody specificity in human brain tissue. Vibratome sections of adult human temporal lobe immunostained for IBA1 (ionized calcium-binding adapter molecule 1, magenta), nuclei dye Hoechst (blue), and antibodies used throughout this study (green): **a**, CD31, platelet endothelial cell adhesion molecule (PECAM-1). **b**, CD45, cluster of differentiation 45/ protein tyrosine phosphatase receptor. **c**, CD163, cluster of differentiation 163/ scavenger receptor cysteine-rich type 1 protein M130. **d**, CX3CR1, Chemokine (C-X3-C) Receptor 1. **e**, MYB, MYB Proto-Oncogene. **f**, P2Y12, Purinergic receptor P2Y G-protein-coupled 12. **g**, PU.1, Hematopoietic transcription factor PU.1. **h**, RUNX1, Runt-related transcription factor 1. **i**, VIMENTIN. Scale bar: 20 μm.

**Supplementary Figure 3.**
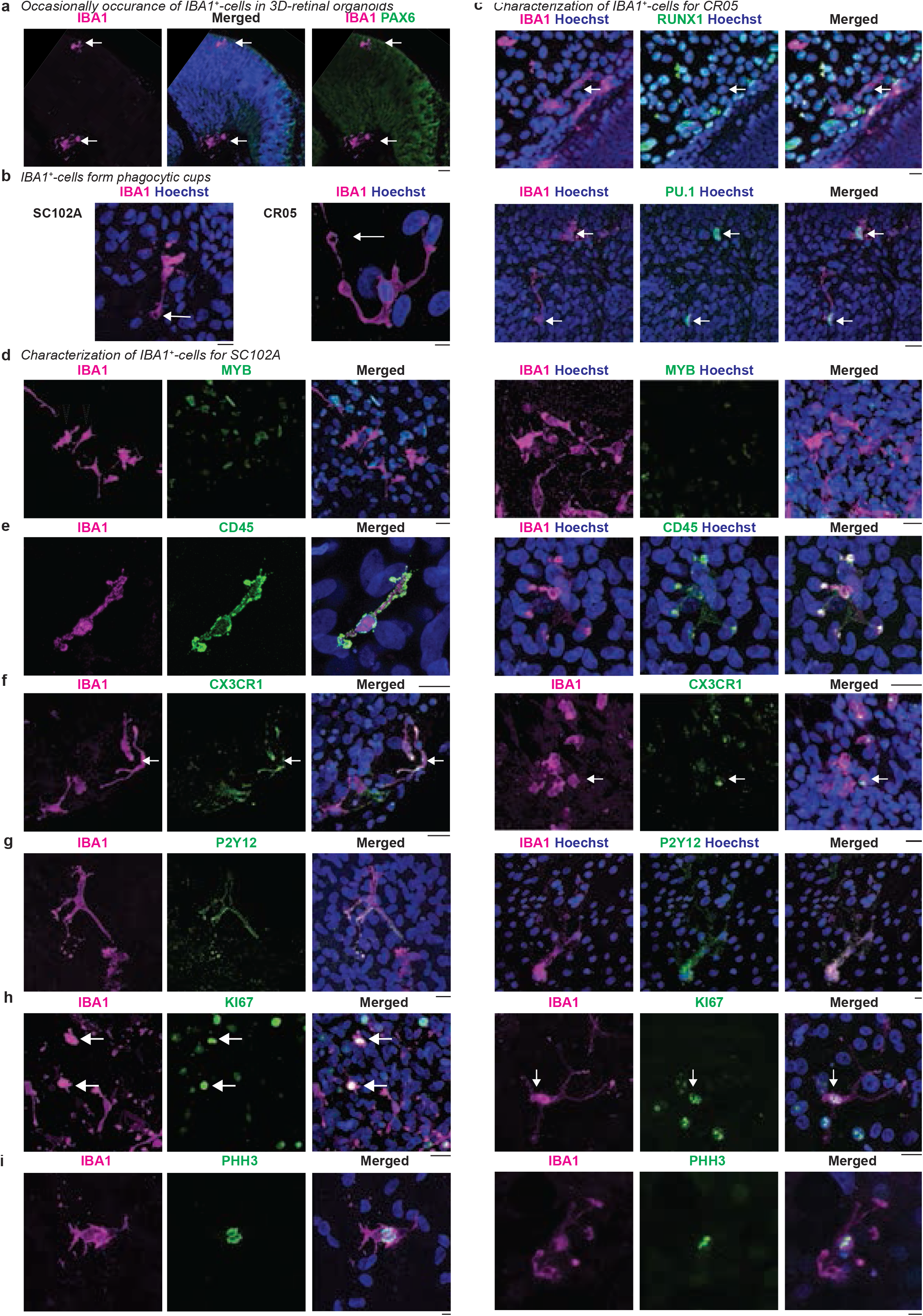
Characterization of IBA1^+^-cells. Immunostaining for IBA1 (ionized calcium-binding adapter molecule 1, magenta) and the nuclei dye Hoechst (blue). **a**, Cryostat section of 3D-retinal organoids differentiated from SC102A at week 9 with focus on retinal-cup. White arrow, examples of overlap. **b**-**i**, Immunostaining in 2.5D culture. Scale bar: 20 μm. **b**, Examples of IBA1^+^-cell forming phagocytic cups (arrow) in 3D structures from SC102A (left) and CR05 (right) at week 8. Scale bar: 10 μm. **c**-**d**, Hematopoietic lineage-specific markers between week 4-5. Immunostaining in green: **c,** RUNX1 (runt-related transcription factor 1), PU.1 (hematopoietic transcription factor PU.1) in CR05. **d**-**i**, SC102A (left) and CR05 (right). **d**, MYB (MYB Proto-Oncogene). **e**-**g**, at week 4-5. **e**, Hematopoietic marker CD45 (cluster of differentiation 45/ protein tyrosine phosphatase receptor). **f**, CX3CR1 (Chemokine (C-X3-C) Receptor 1). **g**, P2Y12 (purinergic receptor P2Y G-protein-coupled 12). **h**-**i**, Proliferation marker between week 8-10 in green: KI67 (**h**) PHH3 (phosphohistone H3, **i**). Scale bar: 20 μm, if not otherwise indicated.

**Supplementary Figure 4.**
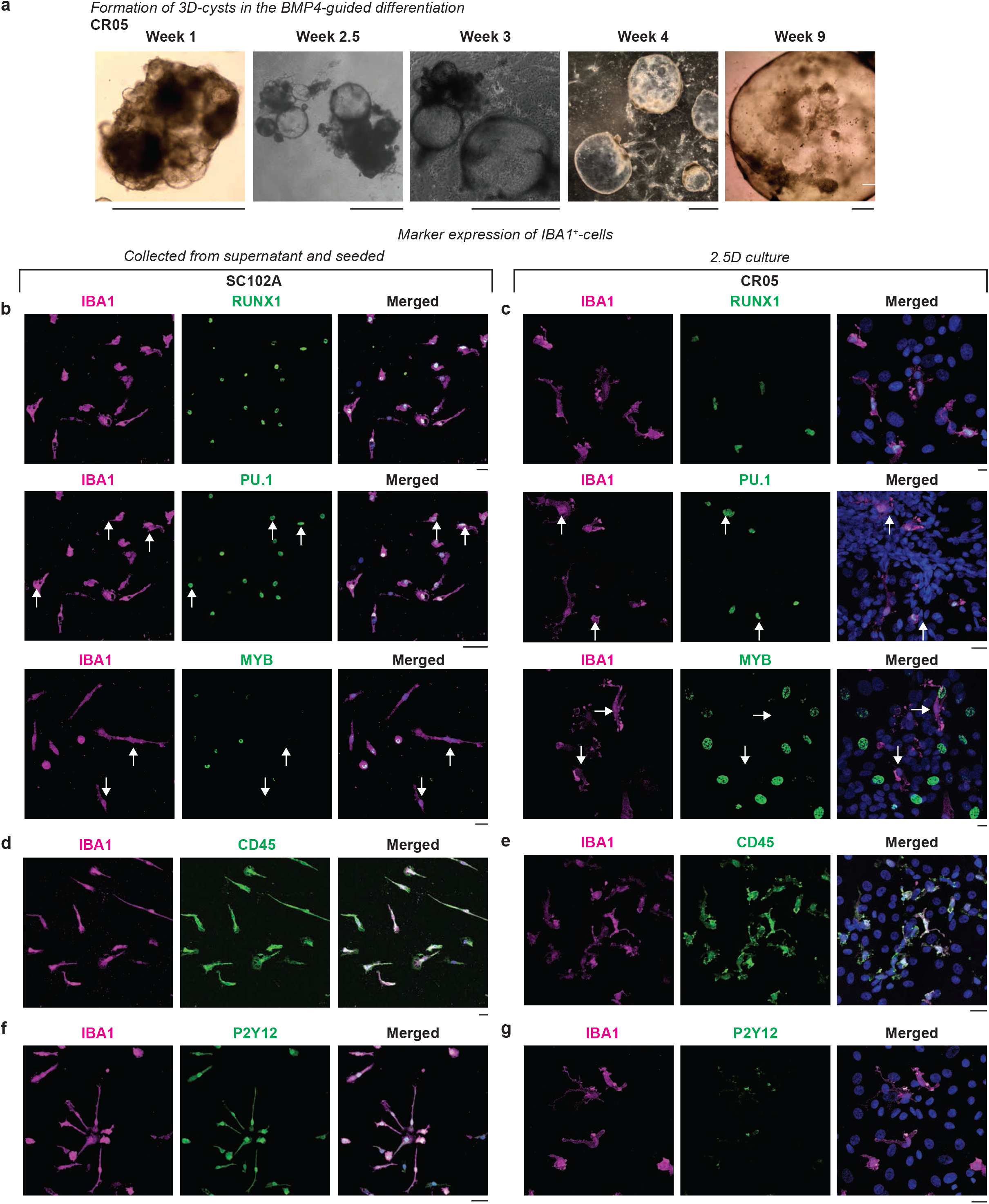
IBA1^+^-cells occur with BMP4-guided differentiation. **a**, Brightfield images of developing 3D-cysts generated with BMP4-guided differentiation for CR05 (week 1-9). Scale bar: 1000 μm. **b**-**i**, Immunostaining for IBA1 (ionized calcium-binding adapter molecule 1, magenta) and nuclei dye Hoechst (blue). **b, d, f**, Cells collected from supernatant of SC102A differentiation and seeded newly at week 6. **c, e, g**, Branched cells within 2.5D culture of CR05 between week 4-6. Scale bar: 20 μm. In green: Immunostaining for **b**-**c**, RUNX1 (runt-related transcription factor 1), PU.1 (hematopoietic transcription factor PU.1), and MYB (MYB Proto-Oncogene). White arrow, examples of non-overlap. **d**-**e**, CD45 (cluster of differentiation 45/ protein tyrosine phosphatase receptor). **f**-**g**, P2Y12 (purinergic receptor P2Y G-protein-coupled 12).

**Supplementary Figure 5.**
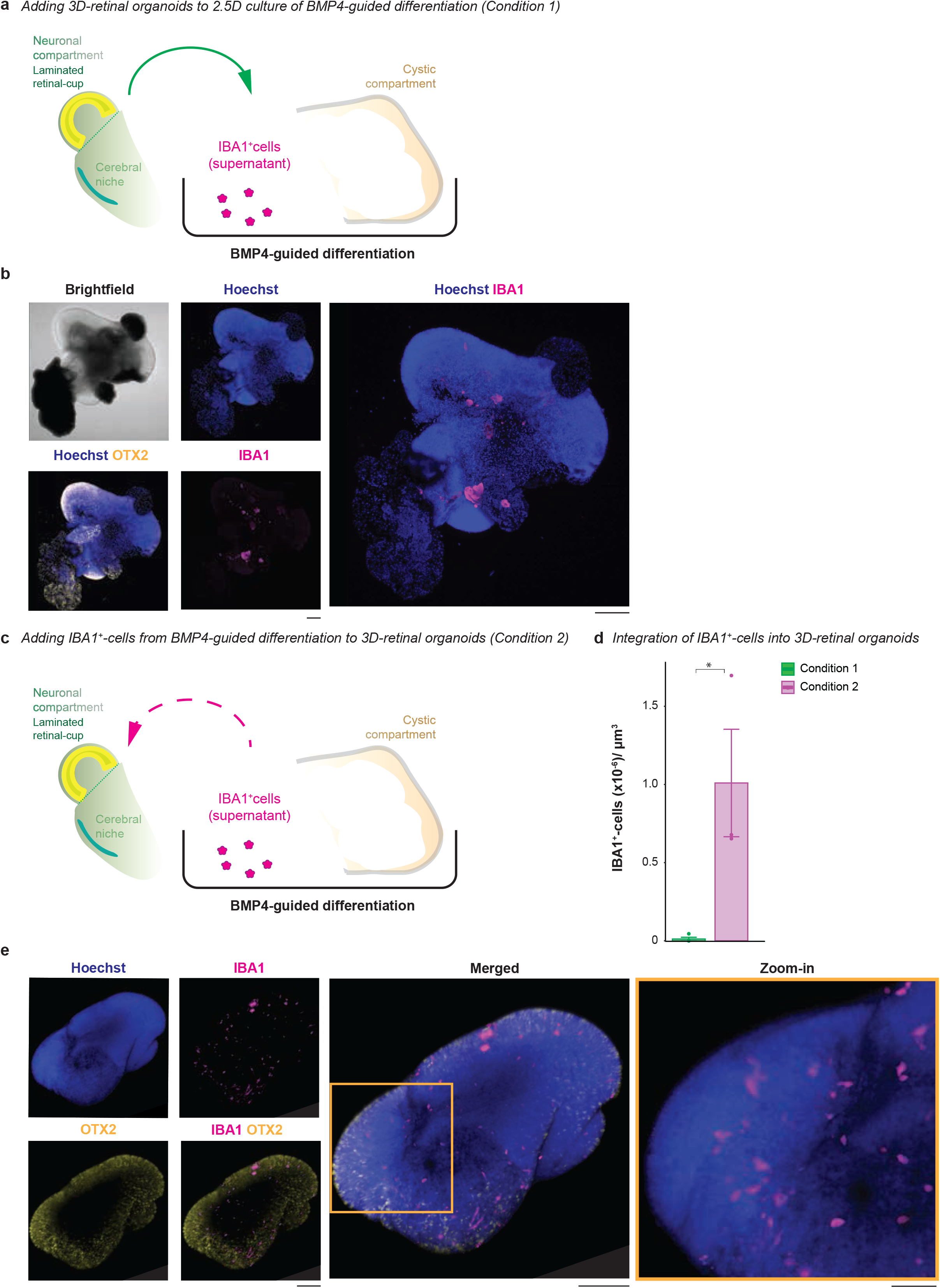
IBA1^+^-cell preference for 3D-cysts *versus* 3D-retinal organoid. **a**, Experimental schematic of adding 3D-retinal organoids to 2.5D culture of BMP4-guided differentiation (condition 1). **b**, Whole mount 3D-retinal organoid from SC102A immunostained for IBA1 (ionized calcium-binding adapter molecule 1, magenta), OTX2 (orthodenticle homeobox 2, yellow), the nuclei-dye Hoechst (blue) and brightfield image at week 18. Scale bar: 100 μm. **c**, Experimental schematic of adding branched cells from the supernatant of BMP4-guided differentiation to 3D-retinal organoids (condition 2). **d**, Bar chart of number of IBA1^+^-cells integrating into the structure and standard error at week 17 after 10 days of adding IBA1^+^-cells. Each dot represents a 3D-retinal organoid. Green, condition 1 related to **a**-**b**. Magenta, condition 2 related to **c, e. e**, Whole mount 3D-retinal organoid immunostaining like in **b**. Wilcox test p-value = 0.0436. *p < 0.05.

